# Probing macromolecular crowding at the lipid membrane interface with genetically-encoded sensors

**DOI:** 10.1101/2023.05.01.538982

**Authors:** Maryna Löwe, Sebastian Hänsch, Eymen Hachani, Lutz Schmitt, Stefanie Weidtkamp-Peters, Alexej Kedrov

## Abstract

Biochemical processes within the living cell occur in a highly crowded environment. The phenomenon of macromolecular crowding is not an exclusive feature of the cytoplasm and can be observed in the densely protein-packed, nonhomogeneous cellular membranes and at the membrane interfaces. Crowding affects diffusional and conformational dynamics of proteins within the lipid bilayer, and modulates the membrane organization. However, the non-invasive quantification of the membrane crowding is not trivial. Here, we developed the genetically-encoded fluorescence-based sensor for probing the macromolecular crowding at the membrane interfaces. Two sensor variants, both composed of fluorescent proteins and a membrane anchor, but differing by the flexible linker domains were characterized *in vitro*, and the procedures for the membrane reconstitution were established. Lateral pressure induced by membrane-tethered synthetic and protein crowders altered the sensors’ conformation, causing increase in the intramolecular Förster’s resonance energy transfer. The effect of protein crowders only weakly correlated with their molecular weight, suggesting that other factors, such as shape and charge play role in the quinary interactions. Upon their expression, the designed sensors were localized to the inner membrane of *E. coli*, and measurements performed in extracted membrane vesicles revealed low level of interfacial crowding. The sensors offer broad opportunities to study interfacial crowding in a complex environment of native membranes, and thus add to the toolbox of methods for studying membrane dynamics and proteostasis.

## Introduction

The interiors of a living cell are recognized as crowded environments, where the concentration of biological macromolecules, predominantly proteins, polynucleotides and their complexes often lays in the range of 300-400 mg/mL (Zimmerman & Trach, 1991; Bohrmann *et al*, 1993; Srere, 1980). This macromolecular crowding decreases the space accessible for biological molecules, thus rendering the “excluded volume effect” (Rivas & Minton, 2018). The excluded volume and the stimulated quinary interactions typically decrease diffusion rates of molecules (Nawrocki *et al*, 2017), affect their conformation and folding (Bai *et al*, 2017; Guseman *et al*, 2018; Berg *et al*, 1999; Kuznetsova *et al*, 2014), and modulate thermodynamic and kinetic properties of biochemical reaction (Minton & Wilf, 1981; Zimmerman & Pheiffer, 1983; Rohwer *et al*, 1998). Although less investigated so far, macromolecular crowding has been also described for the cellular membranes, where the heterogeneous lipid bilayer and ubiquitous integral and peripheral proteins build a complex fluid mosaic structure (Dupuy & Engelman, 2008; Löwe *et al*, 2020). The crowding levels mediated by the membrane proteins, anchored cytoskeleton and eventually polysaccharides are highly specific for cell types and intracellular localization. In red blood cells, proteins occupy 25-30% of the total plasma membrane area (Dupuy & Engelman, 2008), but the protein content may reach 50% within the light-sensitive membrane of the eye rod (Fotiadis *et al*, 2003), and further up to 80% in the densely packed thylakoid membranes (Kirchhoff, 2008; Liu & Scheuring, 2013). This high spatial density of proteins within the lipid bilayer or associated with the membrane interface affects essential cellular processes, including transport across the membrane, cell signalling and energy metabolism, but also the membrane morphology on the meso-scale (Löwe *et al*, 2020).

Despite being an intrinsic property of the cellular membranes, the macromolecular crowding is rarely addressed in molecular studies performed either in native or reconstituted membrane systems. One bottleneck here is quantification of the crowding level and mimicking it appropriately with either proteinaceous or synthetic crowding agent. Previously, a few attempts have been taken to assess the crowding in lipid membranes using non-invasive fluorescence-based approaches. In an early example, crowding-dependent dimerization of the fluorescently- labeled glycophorin A was studied when monitoring changes in Förster’s resonance energy transfer (FRET) (Chen *et al*, 2010). Another approach for the measurement of the interfacial membrane coverage was proposed by the group of Stachowiak and co-workers (Houser *et al*, 2020). The developed synthetic system comprised a polyethylene glycol (PEG) chain anchored at the membrane interface and bearing a donor fluorophore on the free end, and acceptor fluorophores incorporated into the membrane plane. Upon binding of protein crowders to the lipid membrane, rendered steric pressure forced the PEG molecules to elongate and extend over the surface, thus causing decrease in the FRET efficiency. Although promising, the approach may not be applicable to native cellular membranes and *in vivo* experiments.

Here, we describe a genetically-encoded sensory protein that targets the membrane interface and is suitable for measuring the interfacial crowding in synthetic and native membranes. The sensor consists of two fluorescent proteins forming a FRET pair (Boersma *et al*, 2015), which are connected via a flexible linker and a hydrophobic domain. The hydrophobic domain serves as an anchor, so the sensor is stably incorporated into the cellular membrane or synthetic liposomes. The sensor is sterically compressed by the soluble and membrane-coupled crowders, so the associated changes in FRET report on the lateral confinement at the membrane interface. We demonstrate that the crowding induced by either proteins or polymers of varying sizes may be determined using the sensor, and the measurements may be carried out also in native cellular membranes, thus offering a robust approach for crowding analysis in complex environments.

## Results

### Design and expression of the crowding sensors

The primary elements of the FRET-based protein sensor are two fluorescent moieties, such as mCerulean and mCitrine fluorescent proteins, which emission and excitation spectra partially overlap, and a flexible linker, whose structural properties may be altered (Boersma *et al*, 2015; Liu *et al*, 2017). Designing a membrane-associates sensor further required: (i) stable anchoring of the sensor within the lipid bilayer or at the interface of the native and reconstituted membranes; (ii) *cis*-configuration of two fluorescent proteins relative to the membrane plane; and (iii) sufficient flexibility of the intramolecular linkers to allow the crowding-dependent conformational dynamics. A transmembrane helical pair, or *hairpin*, was considered as a suitable membrane anchor, where the fluorescent proteins could be positioned at its N- and C-terminal ends. Firstly, membrane-embedded helical pairs play an important role in membrane protein folding and manifest high stability within the lipid bilayer (Engelman & Steitz, 1981; Kedrov *et al*, 2004; Janovjak *et al*, 2004). Secondly, a helical hairpin would ensure the appropriate topology of the sensor, so the fluorescent proteins would be positioned in proximity to each other at the same side of the membrane.

The recent structure of the membrane-embedded SecYEG translocon of *E. coli* visualized a helical hairpin built of TMHs 1-2 of SecE (Kater *et al*, 2019) (Suppl. Figure 1A). Although being a part of the quaternary complex, the hairpin has minimal contacts with other TMHs of the translocon or within the translocon dimer, and so it forms a stably folded structural unit (Breyton *et al*, 2002). Indeed, the SecE TMH 1-2 hairpin, optionally extended with either a N- or C-terminal soluble domain, was efficiently expressed in *E. coli* as a membrane protein, validating the choice of the potential anchor (Suppl. Figure 1B). Next, the SecE hairpin was cloned into the middle of the soluble crowding sensor (Boersma *et al*, 2015) resulting in two constructs, where the intramolecular linkers either consisted of flexible Gly-Ser-Gly repeats (further referred as (GSG)_6_-SecE) or also contained Glu-Ala-Ala-Ala-Lys repeats forming soluble *α*-helices (*α*H-SecE; Figure 1A). Both sensors were overexpressed in *E. coli* and incorporated into membranes as the full-length proteins, while the degradation products were largely localized to the cytoplasmic fraction (Suppl. Figure 2). Repetitive washes of the membrane fraction, also with either sodium carbonate or urea, which remove loosely attached peripheral proteins, did not affect the localization of the sensor molecules (Figure 1B). Thus, the hydrophobic helical hairpin ensured stable anchoring of both constructs within the membrane.

**Figure 1.**
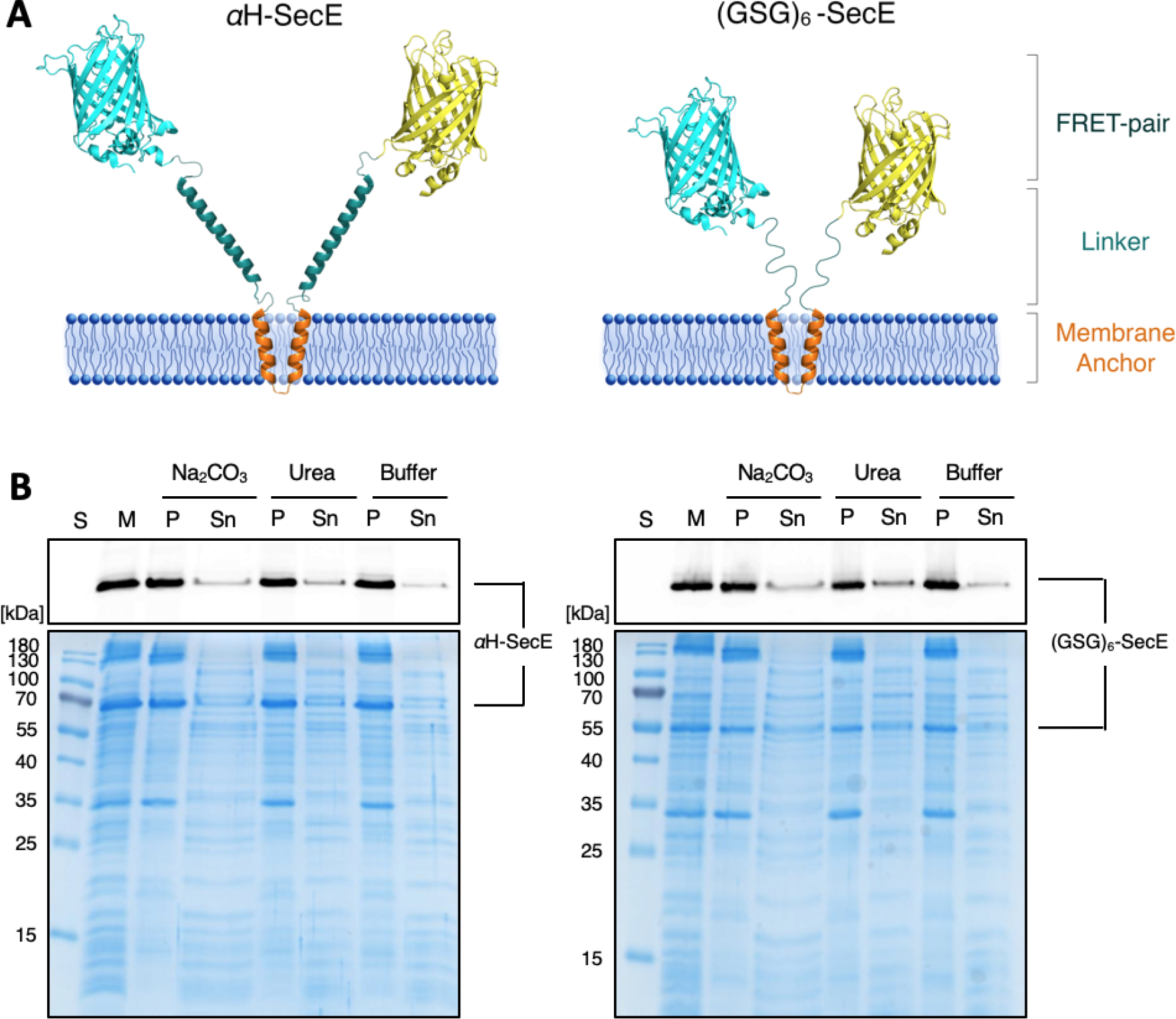
Design and expression of membrane crowding sensors. (A) Schematic representation of the designed sensors. (B) SDS-PAGE of crude membrane extract containing the indicated sensor prior (“M”) and after incubation/washing with either Na_2_CO_3_, urea or the storage buffer. “P” and “Sn”, pellet and supernatant fractions after the incubation, respectively. “S”, PageRuler Prestained Protein ladder. Top: In-gel fluorescence, bottom: Coomassie- stained gels.

To isolate the sensors for further characterization, the membranes were solubilized with 1 % n-dodecyl β-D-maltoside (DDM) and the tagged sensors were purified via the metal affinity and size exclusion chromatography (SEC; Figure 2A and B). The migration of the sensors on SEC was unexpectedly fast for the proteins of ∼70 kDa, but could be potentially explained by the presence of DDM micelle of 76 kDa (Strop & Brunger, 2005), extended protein conformations and/or protein oligomerization. The molecular weights and the oligomeric state of both sensors were analyzed then by SEC coupled to multi-angle light scattering (SEC-MALS; Figure 2B). After subtracting the predicted mass of the DDM micelle, the average molecular weights were 84 ± 3 kDa for *α*H-SecE and 92 ± 1 kDa for (GSG)_6_-SecE sensors. These values exceeded the weights of the monomeric sensors and suggested partial dimerization, which could be induced at the elevated protein concentration of 0.55 mg/mL in the SEC-MALS experiment. To tackle whether the dimerization is dependent on the hydrophobic anchor, we examined a mutant sensor where the anchor domain was substituted with a polar polypeptide. While the calculated molecular mass of the protein is 59 kDa, the apparent mass determined in SEC-MALS experiments ranged from 72 kDa at 0.5 mg/mL to 80 kDa at 3.4 mg/mL (Suppl. Figure 3), and even larger species with the mass to 130 kDa could be resolved. Thus, the oligomerization propensity of the sensors at high concentrations could be related to the constituting fluorescent proteins, but unlikely to have substantial influence at the low levels of the sensor required for the spectroscopy applications.

**Figure 2.**
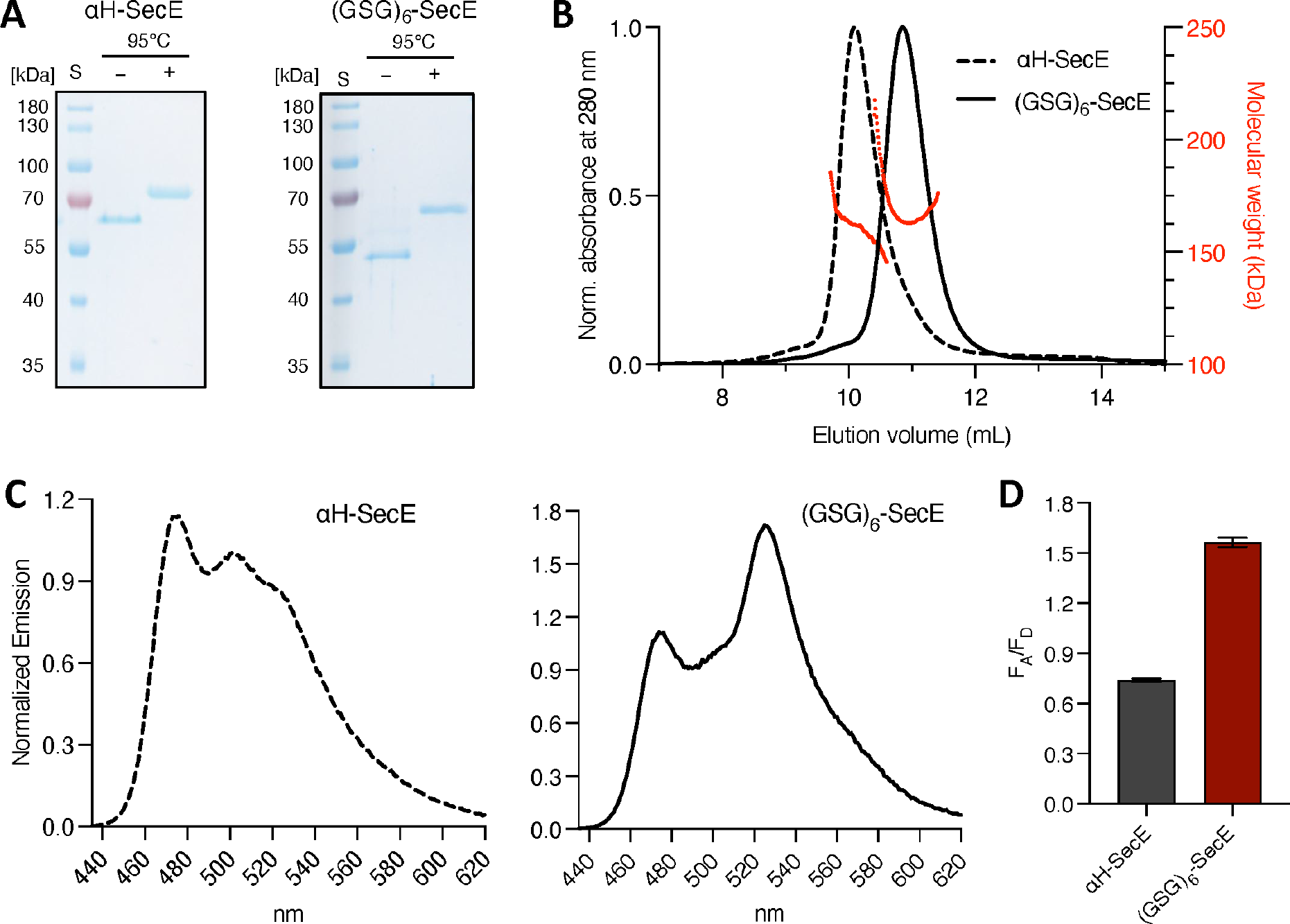
Isolation and characterization of the membrane crowding sensors. (A) SDS- PAGE of purified *α*H-SecE and (GSG)_6_-SecE sensors, with and without thermal denaturation. “S”, PageRuler Prestained Protein ladder. (B) SEC-MALS profiles of the purified sensors constructs and determination of the molar masses. (C) Fluorescence emission spectra of purified and detergent-solubilized sensors (normalized at 500 nm). (D) Calculated FRET efficiency for the detergent-solubilized sensors.

### Spectroscopic characterization of the crowding sensors

The absorbance spectra of purified and detergent-solubilized *α*H-SecE and (GSG)_6_-SecE sensors manifested the specific peaks for mCerulean and mCitrine at 433 and 515 nm, respectively (Suppl. Figure 4), and the difference in the peak intensities correlated with the extinction coefficients of the fluorescent proteins (ε ^433nm^ = 33,000 M^-1^ cm^-1^, ε ^516nm^ = 94,000 M^-1^ cm^-1^). The emission spectra of both sensors (Figure 2C) and the ratio between the acceptor and donor fluorescence at 525 and 475 nm, respectively (further indicated as F_A_/F_D_ ratio), provided the information about the FRET efficiency, and so the sensor conformation. F_A_/F_D_ ratios measured for the detergent-solubilized sensors were 0.74 ± 0.01 for *α*H-SecE, and 1.56 ± 0.03 for (GSG)_6_-SecE (Figure 2D). Thus, the folded helices within the linker domains of *α*H-SecE ensured wider spacing between the fluorescent moieties. Interestingly, the values correlated with those previously measured for soluble sensors (Liu *et al*, 2017): In absence of the membrane anchor the soluble sensors with comparable linker architectures manifested F_A_/F_D_-ratios of 0.55 for the sensor GE (analog of *α*H-SecE) and 1.4 for the sensor G12 (analog of (GSG)_6_-SecE).

The detergent-solubilized sensors were examined for their propensity to respond to crowding upon increasing concentrations of polyethylene glycol (PEG) 6000 in solution (Suppl. Figure 5). PEG is an inert synthetic polymer commonly used as a mimetic crowding agent (Aumiller *et al*, 2014; Kuznetsova *et al*, 2015; Liu *et al*, 2017). The hydrodynamic radius of PEG 6000 is 2.5 nm (Armstrong *et al*, 2004) that can be compared to the dimensions of lysozyme (2.2 nm) or GFP (2.8 nm) (Elowitz *et al*, 1999; Nemzer *et al*, 2013). Increasing PEG 6000 concentration from 0 to 30 % (w/v) led to the substantial increase of the acceptor fluorescence, and so the FRET efficiency for both sensors (Suppl. Figure 5). In the presence of 30% PEG, the F_A_/F_D_ ratio reached 1.85 ± 0.04 for *α*H-SecE and 3.20 ± 0.06 for (GSG)_6_-SecE, suggesting the compression of the flexible sensor molecules under the steric forces. Diluting PEG 6000 from 20% to 10% caused a decrease of F_A_/F_D_-ratios, so both sensors possessed sufficient flexibility to reversibly react to the crowding levels.

### Reconstitution of the sensor into lipid membranes

To characterize the performance of the sensors at the lipid interface, they were reconstituted into liposomes composed of DOPC:DOPG lipids (molar ratio 7:3). Varying the sensor-to-lipid ratio allowed determining the effect of intermolecular FRET between the reconstituted sensors: The F_A_/F_D_ ratios measured in liposomes at the ratios 1:3,000, 1:10,000 and 1:20,000 were comparable with each other, with variations typically within 5 % (Figure 3A). However, when the sensor-to-lipid ratio reached 1:1,000, the FRET efficiency rapidly increased by approx. 20 % for each sensor. Similar concentration dependence was observed for mCerulean-SecE and SecE-mCitrine co-reconstituted in liposomes (Suppl. Figure 6), thus pointing to intermolecular FRET at elevated protein-to-lipid ratios, either due to random contacts or due to clustering of the sensors in the lipid membrane. Based on those insights and the optimal signal-to-noise level, further experiments were conducted at the reconstitution ratio of 1:3,000, where a single sensor molecule would occur on average over 1,000 nm^2^ area of the lipid bilayer (Hills & McGlinchey, 2016; Kamel *et al*, 2022).

**Figure 3.**
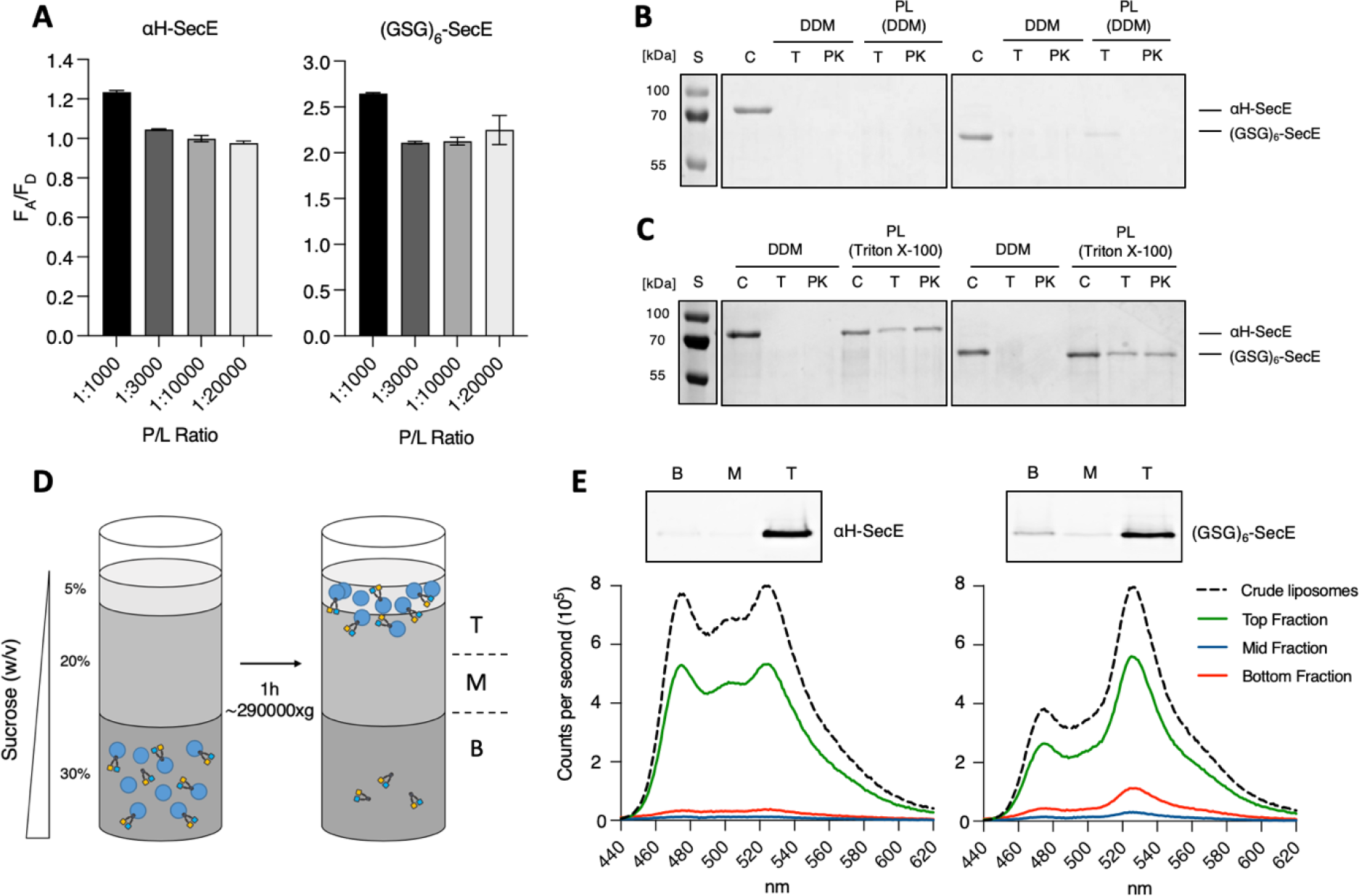
Reconstitution of the crowding sensors in liposomes. (A) FRET efficiencies manifested by the sensors reconstituted at indicated protein-to-lipid (P/L) ratios. (B) Topology determination of the liposome-reconstituted sensors via limited proteolysis by trypsin, “T”, or proteinase K, “PK”. “DDM”, detergent-solubilized sensors, “PL (DDM)”, sensors in proteoliposomes reconstituted using DDM. (C) Same as (B), but using Triton X-100 for the reconstitution. (D) Scheme of the flotation assay. Fractions collected after centrifugation: T (top), M (middle), B (bottom). (E) In-gel fluorescence and fluorescence emission spectra of the sensors in crude liposomes and in fractions collected from the flotation assay.

Next, we examined the topology of the reconstituted sensors, and so, their accessibility to the crowding agents, which could be added in the following steps. The topology was determined based on the sensor susceptibility to trypsin and proteinase K, two proteases with a broad specificity, which could completely degrade the detergent-solubilized sensors (Figure 3B and Suppl. Figure7). For the reconstituted sensors, the proteases can only process the accessible parts of the molecule exposed to the exterior of the liposome, such as the linker domains, and the degradation may be monitored via SDS-PAGE. Upon the proteolytic treatment of the liposome-anchored sensors, the bands for the full-size proteins disappear for all samples, with an exception for the (GSG)_6_-SecE FRET sensor treated with trypsin. Here, the digest has not been accomplished completely. While the lysine-containing *α*-helices in the *α*H-SecE construct offer multiple cleavage sites, those not present within the linkers of (GSG)_6_-SecE, resulting in the partial proteolysis. The results implied that the majority of the liposome-reconstituted sensors have the outward-facing orientation. As a control, we additionally performed the same experiment with proteoliposomes where the lipids were treated with 0.5 % Triton X100 detergent prior to reconstitution (Figure 3C). Under these conditions, the liposomes are rather solubilized than swelled (Suppl. Figure 8), which favors dual, stochastically-driven orientation of the sensor in the liposomes (Geertsma *et al*, 2008; Niroomand *et al*, 2016). The pattern of the protected bands observed on SDS-PAGE after the protease treatment suggested that 30 to 50 % of sensors indeed acquired the inward-facing orientation (Figure 3C).

Notably, even at low sensor-to-lipid ratios F_A_/F_D_ values in liposomes was by 25-30% higher than those recorded for the detergent-solubilized sensors (Figure 2D). To examine whether the increased FRET signal is caused by the off-pathway aggregation, we analyzed the sensor reconstitution efficiency. Once loaded into the sucrose gradient, the liposomes could float to the top due to the density difference between the aqueous interior and the external solution (Figure 3D). Only reconstituted sensors were able to co-migrate with the liposomes, while the non-reconstituted and aggregated proteins remained at the bottom of the gradient. The analysis of the collected fractions by SDS-PAGE showed that both sensor variants predominantly appeared in the top fraction (Figure 3E). The reconstitution efficiency reached 96 % for *α*H-SecE and 84 % for (GSG)_6_-SecE sensors. The proteins remaining in the minor bottom fraction manifested a high FRET efficiency, as F_A_/F_D_ ratio reached 2.59 ± 0.10 for (GSG)_6_-SecE sensor (not determined for αH-SecE due to the low concentration in the bottom fraction), as could be expected from the clustered/aggregated molecules. The FRET efficiency of (GSG)_6_-SecE sensor in the top fraction was 2.08 ± 0.01, that matched closely the value measured for the crude reconstituted sensor, 2.13 ± 0.02. For *α*H-SecE sensor prior and after the flotation assay the values were nearly identical, 1.03 ± 0.02 and 1.012 ± 0.004, respectively (Figure 3D). Thus, we concluded that the sensors were successfully reconstituted into liposomes, and the resulting relatively high FRET efficiencies were due to altered conformations of the sensors in presence of the proximate lipid interface.

### Sensitivity of the reconstituted sensor constructs to crowders

Increased FRET efficiency for the liposome-reconstituted sensors suggested that the proteins acquired more compact conformations at the membrane interface. We questioned whether the sensors remained sufficiently dynamic to respond to the changes in the proximate crowding. To test that, soluble PEG 2000 and 6000 were added to the proteoliposome suspension. Upon increasing PEG 6000 concentration up to 30 % (v/v), FRET efficiency increased up to 3.00 ± 0.03 for *α*H-SecE (increase by 175 %, Figures 4A to C) and to 3.78 ± 0.04 for (GSG)_6_-SecE (increase by 87 %; Figure 4D to E). Thus, despite the constraints set by the membrane interface, both sensors were responsive to the surrounding crowding levels. In the next step, the performance of the sensors was studied in the presence of the interfacial polymer crowding. For this purpose, PEG-grafted lipids (DOPE-PEG 2000) were incorporated into the liposomes. PEG 2000 at the interface should render the lateral pressure (Marsh *et al*, 2003), which may cause compression of the membrane-anchored sensors (Figure 5A). Both sensors responded to the changes in the interfacial crowding, as the FRET efficiency increased nearly linearly with increasing concentration of DOPE-PEG 2000 (Figure 5B to E). In presence of 10 mol % DOPE-PEG 2000, the FRET efficiency reached 1.20 ± 0.02 for the *α*H-SecE (increase by 16 %), and 2.86 ± 0.17 for the (GSG)_6_-SecE construct (increase by 33 %).

**Figure 4.**
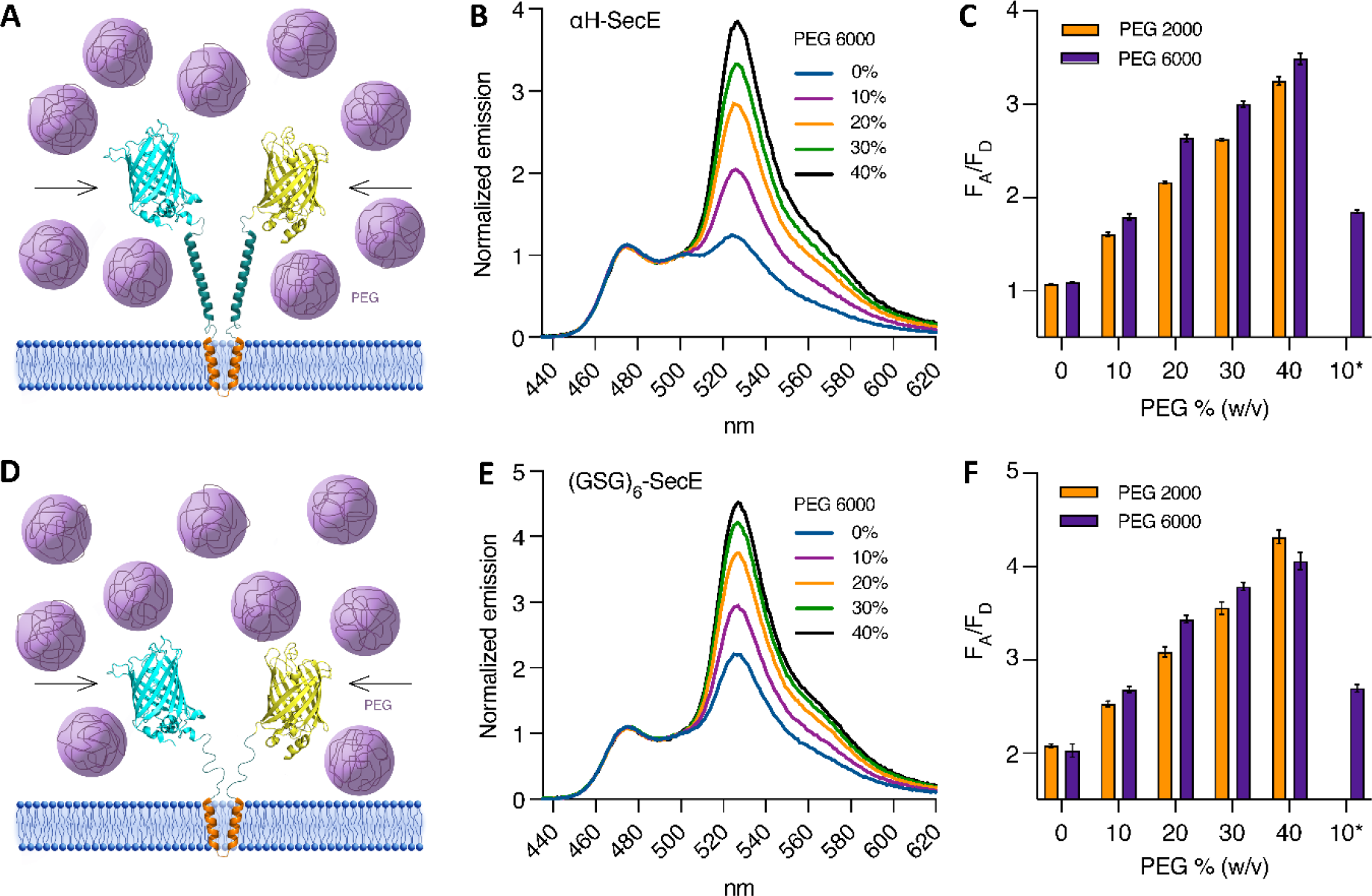
Sensitivity of the membrane-anchored sensors to soluble crowders. (A) Scheme of the reconstituted *α*H-SecE sensor in presence of PEG molecules in solution. (B) Fluorescence emission spectra of *α*H-SecE in presence of PEG 6000 at indicated concentrations (w/v). The spectra are normalized at 500 nm. (C) FRET efficiencies of *α*H-SecE in presence of PEG 6000 or PEG 2000 (mean ± SD, n = 2). Samples “10%*” correspond to two-fold dilution of 20% PEG 6000 for testing the reversibility of the sensor compaction. (D-F) Same as (A-C), for the liposome-reconstituted (GSG)_6_-SecE sensor.

**Figure 5.**
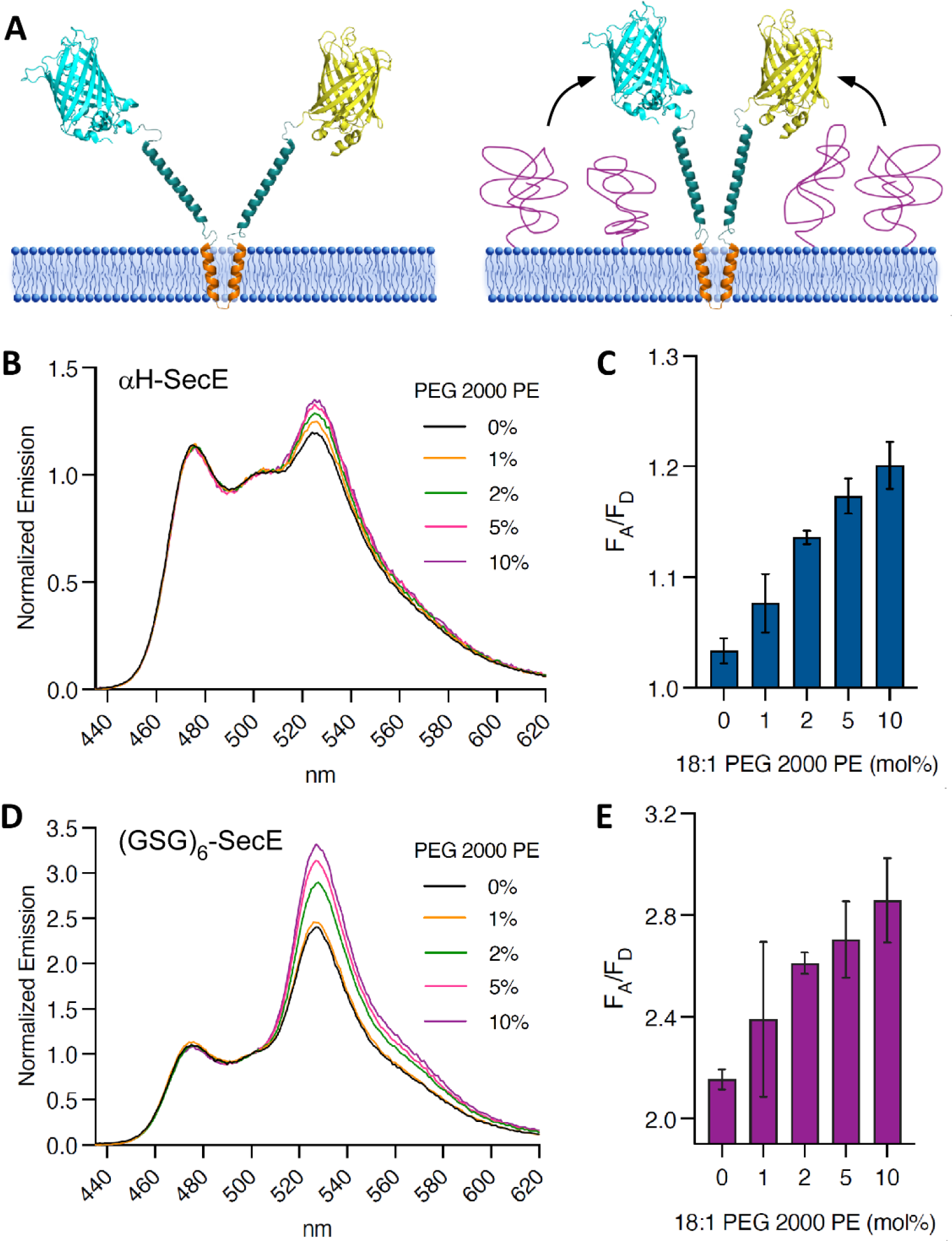
Sensitivity of the membrane-anchored sensors to interfacial polymer crowding. (A) Scheme of the reconstituted *α*H-SecE sensor upon compaction induced by a polymer at the membrane interface. (B and C) Fluorescence emission spectra and corresponding FRET efficiencies (mean ± SD; n = 2) of *α*H-SecE sensor in presence of DOPE-PEG 2000 lipids at indicated concentrations (mol %). (D-E) Same as (B-C), for the reconstituted (GSG)_6_-SecE sensor.

To generate native-like protein-based crowding, proteins of choice could be specifically anchored at the membrane interface via either Ni^2+^-NTA:histidine or biotin:streptavidin coupling. To ensure anchoring of various poly-histidine-tagged proteins, 18:1 DGS-NTA lipids were incorporated into liposomes, while the tag-less sensors were employed for the reconstitution. The following poly-histidine-tagged proteins were used then as crowders: monomeric streptavidin (mSA; molecular mass 15.5 kDa) (Demonte *et al*, 2014), SecB chaperone (monomer size 20.3 kDa), and SecA ATPase with either N- or C-terminal polyhistidine-tags (SecA^N^ and SecA^C^, monomer size ∼100 kDa) (Figure 6A and 6B). Among those, SecB forms a stable tetramer, thus reaching approx. 80 kDa mass (Smith *et al*, 1996), while SecA may exist both in monomeric and dimeric forms, but predominantly monomeric once it is bound to the membrane (Roussel & White, 2020). Various amounts of the crowders were incubated with proteoliposomes to achieve either partial or complete coverage of the surface-exposed Ni^2+^-NTA groups (Figure 6C) (Raghunath & Dyer, 2019). All the examined protein crowders induced the concentration-dependent response of the *α*H-SecE sensor, but the measured FRET efficiencies were protein-specific. Thus, titration of the ATPase SecA, the largest examined crowder with either N- or C-terminal anchoring tag, induced a rapid increase in the F_A_/F_D_ ratio followed by a plateau, indicating saturation of the liposome surface with the bound crowder. Notably, different FRET efficiencies were achieved when using either N- or C-terminally-tagged SecA variants (SecA^N^ and SecA^C^), with the maximal increase of 14 % and 8 %, respectively. Strikingly, the relatively small protein mSA induced an equal increase in the FRET efficiency as the N-terminally bound SecA^N^ ATPase, while the tetrameric SecB caused the minimal change in the FRET signal (Figure 6C). Thus, the molecular weight of a crowder was not the decisive factor for the intensity of the sensor response. At the end of the experiment, the proteoliposomes were incubated with imidazole to dissociate the crowders from the surface, and the FRET efficiency dropped to the initial crowder-free values. Disrupting the proteoliposomes with 1 % DDM caused further decrease of the F_A_/F_D_ ratio to 0.74, matching the value measured for the detergent-isolated sensor. The non-tagged streptavidin variant Strep^D4^ of 60 kDa served as a negative control, which did not affect the fluorescence, and so the sensor conformation (Howarth *et al*, 2006).

**Figure 6.**
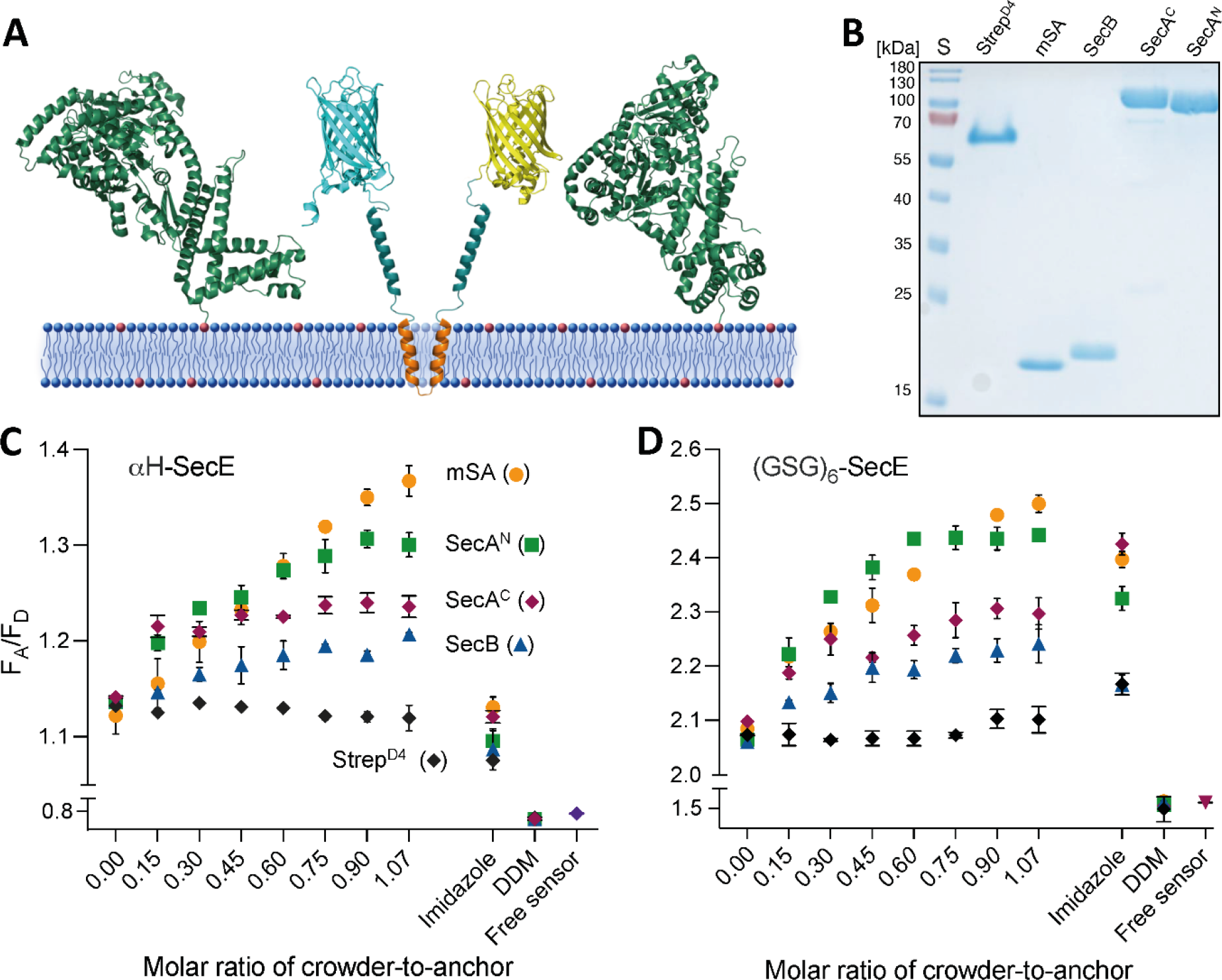
Sensitivity of the membrane-anchored sensors to interfacial protein crowding. (A) Scheme of the reconstituted *α*H-SecE sensor in presence of protein crowders, e.g. SecA (green) anchored at the membrane interface via specific protein:lipid contact sites (red dots). (B) SDS-PAGE of purified proteins applied as crowders. “S”, PageRuler Prestained Protein ladder. (C) FRET efficiencies of the sensors in presence of increasing concentrations of the protein crowders. “Imidazole”, FRET signal after adding 300 mM imidazole to detach the crowders. “DDM”, FRET signal after adding detergent to extract the sensor from the membrane. “Free sensor”, FRET signal of the sensor prior to the liposome reconstitution.

Qualitatively similar results were obtained when employing proteoliposomes with (GSG)_6_-SecE sensor, as the N-terminally anchored SecA^N^ and mSA induced the most prominent increase in FRET (Figure 6D). However, addition of imidazole could only partially reduce the FRET signal of the sensor, and not for all tested crowders. Notably, the signal even increased for the Strep^D4^ protein that served as a negative control. Since the reversibility of the sensor dynamics in response to changes in crowding was previously confirmed (Figure 4F), we suspect that the elevated imidazole concentration caused unpredicted conformational rearrangements within the flexible linkers, not related to the crowding *per se*. Nevertheless, excess of the detergent added to proteoliposomes triggered the decay in the FRET efficiency to the level of membrane- free sensor (Figure 6D).

In an alternative approach, the liposomes with αH-SecE sensor were supplemented with 18:1 biotinyl cap PE lipids, so the crowder proteins could be deposited at the lipid membrane interface via biotin:streptavidin coupling (Suppl. Figure 9). Here, mSA played the role of the crowding agent, and its effect on the sensor conformation could be compared for two binding modes, i.e. via NTA and biotin anchoring, as the protein contained a poly-histidine tag (Figure 6C). For the biotin-functionalized liposome containing αH-SecE sensors, continuous increase in the FRET efficiency was observed upon titrating mSA suggesting compression of the sensor (Suppl. Figure 10). At the highest examined mSA concentration, the F_A_/F_D_ reached 1.37 ± 0.03, which indicates increase of the FRET efficiency by 22%, and the response of the sensor to the increasing mSA concentration was similar between biotin and Ni^2+^-NTA surface anchors.

Finally, we examined whether αH-SecE or (GSG)_6_-SecE are responsive to the crowding within the lipid bilayer. For this purpose, the sensors were reconstituted into liposomes (protein-to- lipid ratio 1:3,000) in presence of the membrane protein complex SecYEG (Suppl. Figure 1 and 11A). *E. coli* SecYEG consists of 15 TMHs connected by relatively short loops, and it lacks large extramembrane domains, so the protein should not render substantial interfacial crowding. Indeed, even at the molar ratio of SecYEG to lipids of 1:300 that corresponds to mass ratio of 1:3 neither of the crowding sensors manifested higher FRET efficiency (Suppl. Figure 11 B). The observation does match the initial intuitive prediction, but it also suggests that the crowding within the membrane does not induce clustering of the sensors, that otherwise would result in high inter-molecular FRET.

### Crowding analysis in cellular membranes

The broad interest in genetically-encoded sensors arises from the opportunity to probe the conditions within the native cellular environments. Characterization of the crowding sensors in synthetic membranes provided above demonstrates their fitness for the proposed task, and we further set out to employ them for measuring the interfacial crowding in a physiologically relevant environment, the inner membrane of *E. coli.* As unambiguous analysis in the living cell would be complicated at this stage by the intrinsically high crowding in the cytoplasm, we pursued measurements in isolated bacterial membranes.

While the low density of the sensors, and so minimal intermolecular FRET in model liposomes could be achieved by adjusting the protein:lipid ratio upon the membrane assembly, the density of the sensors in the cellular membrane should be controlled by tuning their expression level. For this purpose, expression of both *α*H-SecE and (GSG)_6_-SecE sensors was carried out using a tightly regulated arabinose-inducible promoter. To validate the membrane localization of the expressed sensors, *E. coli* host cells were imaged by super-resolution structured illumination microscopy (SR-SIM) (Figure 7A). For both sensors, fluorescence signal of the acceptor fluorophore mCitrine was observed along the contour of individual bacteria verifying the localization of the proteins at the membrane. Though the expression level was notably higher for *α*H-SecE, the fluorescence signal of both variants was homogeneously distributed over the cell surface without cluster formation or accumulation at the poles.

**Figure 7.**
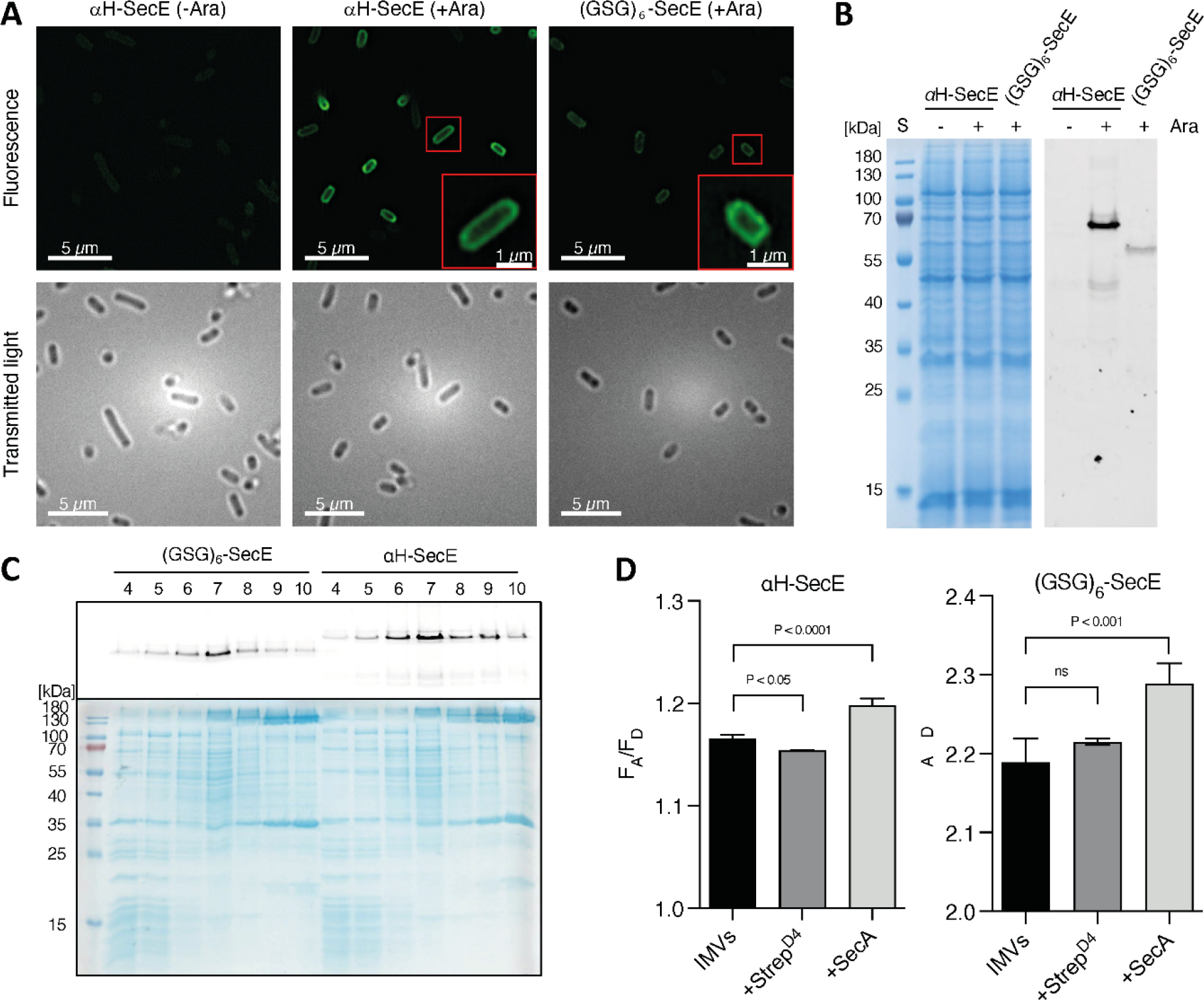
**Crowding sensors in cellular membranes**. (A) Super-resolution fluorescence (top) and corresponding transmitted microscopy images (bottom) of the *E. coli* cells expressing *α*H- SecE and (GSG)_6_-SecE FRET sensors. Uninduced cells bearing *α*H-SecE expression plasmid served as control (“-Ara”). (B) SDS-PAGE of total cell protein extracts with and without sensor overexpression. Left: Coomassie stained gel; right: in-gel fluorescence. (C) SDS-PAGE of sucrose density gradient fractions to separate inner and outer bacterial membranes. Top: in- gel fluorescence; bottom: Coomassie stained gel. Fractions 4 and 5 demonstrate the characteristic pattern of ribosomal proteins, followed by IMVs (fractions 6 to 8). (D) FRET efficiencies of the crowding sensors recorded in IMVs and in presence of either Step^D4^ or membrane-binding SecA (mean ± SD, n= 3)

The presence of both sensor in the membrane was further confirmed by SDS-PAGE in-gel fluorescence of the crude membrane extracts, and the fluorescence intensities correlated with SR-SIM results (Figure 7B). The inner and outer membrane vesicles (IMVs/OMVs) were then separated from each other by sucrose density gradient, and the sensors were predominantly found in the IMV-containing fractions (Figure 7C). To estimate the relative amount of the expressed sensors, we determined the total membrane protein concentration by a colorimetric assay, and the concentration of the sensor by SDS-PAGE in gel-fluorescence, where the independently purified sensor served for the signal calibration (Suppl. Figure 12). *α*H-SecE sensor constituted 3.3 % of the total membrane protein mass, and the fraction of weakly expressed (GSG)_6_-SecE did not exceed 2 % of the total protein content.

With that relatively low abundance of the sensors, and absence of the aggregation clusters in the cells (Figure 7A), we assumed that the intermolecular FRET would not substantially contribute to the fluorescence read-out, and the FRET signal could be related to the crowding- dependent conformations of the sensors. For the extracted IMVs, the F_A_/F_D_-ratio was 1.17 ± 0.01 and 2.19 ± 0.03 for *α*H-SecE and (GSG)_6_-SecE sensors, respectively (Figure 7D), being within the value range measured previously for the synthetic membranes, either in the presence of PEG or proteinaceous crowders (Figures 5 and 6), and corresponding to the low crowding levels. Addition of Strep^D4^ did not influence the sensor conformation, as the F_A_/F_D_- ratios were not affected (1.15 ± 0.01 for *α*H-SecE and 2.21 ± 0.01 for (GSG)_6_-SecE; Figure 7D), the protein was not expected to interact with the membrane surface. To induce the interfacial crowding, we employed the ATPase SecA, as the protein contains an amphipathic N-terminal helix essential for docking SecA at the membrane interface (Kamel *et al*, 2022). Addition of SecA had a weak, but reproducible effect on both sensors, as the FRET efficiencies increased to 1.20 ± 0.01 and 2.29 ± 0.03 for *α*H-SecE and (GSG)_6_-SecE, respectively (Figure 7D).

## Discussion

While the effects of macromolecular crowding on biological membranes are ubiquitous and diverse (Löwe *et al*, 2020; Guigas & Weiss, 2016), the methods to study the crowding in living cells and reconstituted systems are currently limited (Chen *et al*, 2010; Houser *et al*, 2020). In this work, we designed and characterized first genetically encoded FRET-based sensors for the quantification of the crowding at the membrane interfaces and showed that a straightforward reconstitution into model membranes renders the sensors suitable for the assigned task. The difference in the structure of the sensors’ linker domains, i.e. flexible Gly-Ser-Gly repeats vs. folded α-helical domains, had a clear impact on the fluorescence read-out, and so the sensor conformations, in agreement with the earlier study (Liu *et al*, 2017). Both in the detergent micelles and at the membrane interfaces, the FRET efficiency, and so the distance between the fluorescent proteins, was substantially higher for (GSG)_6_-SecE sensor than for *α*H-SecE. Thus, the *α*-helices within the linkers of *α*H-SecE served as spacers within the FRET pair in the absence of crowders, while the unstructured Gly-Ser-Gly repeats rendered a rather compact initial conformation. Nevertheless, both sensors were sufficiently dynamic to respond to the changes in macromolecular crowding induced with either soluble or membrane-associated molecules. Strikingly, while soluble PEG molecules manifested an immense effect, as the FRET efficiencies of the sensors increased 2-3-fold in the presence of 40 % PEG 2000, the same crowder caused rather moderate response when being anchored to the membrane: At the maximal abundance of 10 mol % of DOPE-PEG 2000, the increase in the FRET efficiency was limited to 16 % for *α*H-SecE and 33 % for (GSG)_6_-SecE sensor. Here, the conformational dynamics of the PEG chains may play a role, as the polymer undergoes an entropy-induced elongation, known as “mushroom-to-brush” transition (Marsh *et al*, 2003) when present at 2-3 mol % which may reduce the entropic pressure on the sensor.

Interested in the perspective to measure physiological crowding in cellular membranes, we analyzed the performance of both sensors in presence of protein crowders. For all tested crowders anchored at the functionalized liposomes, both sensors manifested elevated FRET signal upon increasing the crowders abundance. Notably though, the increase in FRET efficiency did not correlate with the molecular sizes of the crowders, as the small protein mSA (16 kDa) and the large motor protein SecA^N^ (∼100 kDa) triggered comparable responses. As the FRET signal commonly reached saturation within the probed crowders concentration range, incomplete binding could be ruled out. Other factors may be the geometry of the crowder binding, as implied by two SecA variants anchored via either N- or C-terminal end, and the tetrameric SecB protein that may acquire planar orientation at the membrane surface when building three or four His:Ni^2+^-NTA contacts. Complementary, the shape and surface charges of the crowders may play roles in quinary interactions with the sensor molecules, so their effect may go beyond the excluded volume (Sarkar *et al*, 2014; Guseman *et al*, 2018; Kuznetsova *et al*, 2015). Determining the complex interactions of various crowders with the sensors is a task for further analysis, where experimental approaches may be combined with computational modelling.

Shown ability of the sensors to target and insert into cellular membranes, together with their functionality within the native membrane vesicles implies applications of the sensors to study membrane proteostasis *in vivo*. Once established in eukaryotic cells, crowding levels may be measured within distinct cellular compartments, and modification of the membrane anchor, i.e. size and hydrophobicity may be used for targeting the sensors to specific organelles or the membrane nanodomains (Sharpe *et al*, 2010; Sezgin *et al*, 2017). Temporarily-resolved experiments may reveal changes in the crowding levels, e.g. due to protein over-expression, membrane stress and cell ageing (Mouton *et al*, 2020; Karagöz *et al*, 2019), and further applied to study the density and dynamics of the cell surface glycocalyx or bacterial lipopolysaccharides. However, both sensors evaluated here demonstrated the prominent response to the crowding in solution proximate to the membrane interface. Although this effect may be beneficial for particular studies, e.g. dynamics of the actin cytoskeleton or assembly of macromolecular condensates proximate to the membrane (Bokvist & Grobner, 2007; Wang *et al*, 2023), uncoupling the sensor dynamics from the solvent conditions is essential to examine exclusively the membrane crowding. Here, further design and optimization of the linker domain architecture is required, that also determines the dynamic range of the sensors, and so the achievable resolution in crowding measurements. Structured domains, such as α-helices in αH-SecE sensor, appear more suitable for design and controlled modifications. Here, introducing amphipathic helices may be a potent strategy, as their crowding-sensitive interactions with the membrane may be employed for switching the sensor conformations (Prévost *et al*, 2018), while variations in the length, charge distribution and flanking elements will serve for further fine-tuning.

Studying organization and dynamics of cellular membranes in a non-invasive manner remains a great challenge in biology, but the recent technical developments, first of all in advanced fluorescence microscopy and membrane-specific probes are providing new tools and opportunities (Sezgin, 2017; Collot *et al*, 2022). We envision that the protein-based sensors for crowding in cellular membranes will be a valuable add-on for characterizing the environment of the cell membrane interfaces, and will also find their applications in crowding analysis in reconstituted systems.

## Materials and Methods

### SecE-FRET-sensor expression and purification from bacterial inner membranes

Gene fragments encoding for TMHs 1-2 of SecE *E. coli* were introduced into the plasmid pRSET-A-FRET (Boersma *et al*, 2015) via Gibson assembly (New England Biolabs), so the encoded membrane anchor substituted the flexible linker between the mCerulean and mCitrine. Additionally, a cleavage site for 3C protease (sequence LEVLFQGPG) was added to each construct after the N-terminal hexa-histidine tag. A soluble sensor contained a polypeptide of 14 amino acids (AHIVMVDAYKPTK) (Zakeri *et al*, 2012) instead of the anchor domain. Cloning results were validated by sequencing analysis (Eurofins Genomics). Resulting plasmids containing genes for (GSG)_6_-SecE and *α*H-SecE sensors were transferred into the *E. coli* C43(DE3) strain. For the protein over-expression, the cultures were grown at 30°C in LB medium (10 g/L tryptone, 10/L g NaCl and 5 g/L yeast extract) supplemented with 100 µg/mL ampicillin till OD_600_ of 0.6 was reached. The expression of the sensors was induced with 0.1 mM IPTG and carried out overnight at 25°C (Boersma *et al*, 2015). For tunable expression of sensors, the constructs were re-cloned into pBAD_His_ vector, and expression was induced with 0.001% L-arabinose. Expression of mCerulean-SecE (pBAD-based vector) and SecE- mCitrine (pRSET-A) was performed using the same protocol.

The cells were harvested by centrifugation at 5000x*g* for 15 min (SLC-6000, Thermo Fisher/Sorvall), resuspended in 20 mM NaH_2_PO_4_/Na_2_HPO_4_ pH 7.4 and 100 mM NaCl supplemented with 0.1 mM PMSF and lysed by Microfluidizer (M-110P, Microfluidics Corp). Cell debris was removed by subsequent centrifugation at 12000x*g* for 15 min (SS34, Thermo Fisher/Sorvall). The membrane fraction was collected by centrifugation for 45 min at 235000x*g* (45 Ti rotor, Beckman Coulter). The pellet was resuspended in 20 mM NaH_2_PO_4_/Na_2_HPO_4_ pH 7.4, 100 mM NaCl, 5% glycerol and 0.1 mM PMSF. Further, the membranes were solubilized in 1 % DDM (Glycon Biochemicals GmbH), 50 mM NaH_2_PO_4_/Na_2_HPO_4_, 500 mM NaCl, 200 µM TCEP and 0.2 mM PMSF. The proteins were purified via metal ion affinity chromatography (IMAC). The solubilized material was loaded on the Ni^2+^-NTA-agarose resin (either QIAGEN or Macherey-Nagel) and the resin was washed with 50 mM NaH_2_PO_4_/Na_2_HPO_4_ pH 8.0, 300 mM NaCl, 0.1 % DDM and 20 mM imidazole. The proteins were eluted with 50 mM NaH_2_PO_4_/Na_2_HPO_4_ pH 8.0, 300 mM NaCl, 0.1 % DDM and 250 mM imidazole. The elution fraction was loaded on the Superdex 200 Increase GL 10/300 column (Cytiva) in 10 mM NaH_2_PO_4_/Na_2_HPO_4_ pH 7.4, 50 mM NaCl and 0.05 % DDM. Peak elution fractions of SEC were pooled, aliquoted and stored at -80°C. The expression of the sensor and each purification stage were controlled via SDS-PAGE, followed by in-gel fluorescence imaging and Coomassie staining (Quick Coomassie^®^ Stain, SERVA). To remove the N-terminal tag, 3С protease was added to the IMAC resin-bound sensors after washing steps and incubated for 2 h. Afterwards, the released protein was eluted with the wash buffer followed by SEC, as described above. For the spectrophotometric analysis, the following extinction coefficients were used to calculate the concentration of fluorescent proteins, and the total protein concentration: mCerulean3 of *ε_433_ =* 33000 M^-1^*cm^-1^, mCitrine of *ε_516_ =* 94000 M^-1^*cm^-1^ (Lambert, 2019). Both sensors, which differ only by the linker sequence, had the extinction coefficient *ε_280_ =* 56520 M^-1^*cm^-1^. The calculated molar ratio of individual fluorescent proteins to the sensor concentration provided an estimate for the folding efficiency. mCerulean and mCitrine of (GSG)_6_-SecE were folded with the efficiency of 61% ± 15% and 73% ± 7%, respectively (three independent expression/isolation experiments). Within the *α*H-SecE sensor, the folding efficiency of the fluorescent domains reached 78% ± 9% and 87% ± 1%, respectively (n=3), suggesting more efficient folding within the construct with the elongated and structured linkers.

### Sensor expression for measurements *in vesicula*

The protein expression using pBAD-based plasmids was conducted as described above, using 0.001% L-arabinose (67 µM) as the inducer. The isolated crude membrane extract was loaded on the continuous 20-70 % sucrose density gradient in 20 mM NaH_2_PO_4_/Na_2_HPO_4_ pH 7.4 and 100 mM NaCl prepared by the Gradient Station (Biocomp) and centrifuged for 16 h at 30.000 rpm (rotor SW 40 Ti, Beckman Coulter). The gradients were collected with the Gradient Station, and the fractions were analyzed on SDS-PAGE. Selected fractions were pooled together, diluted 5-fold with 20 mM NaH_2_PO_4_/Na_2_HPO_4_ and 100 mM NaCl, and pelleted via centrifugation for 45 min at 235,000 g (45 Ti rotor, Beckman Coulter) to remove sucrose. The pellet was resuspended in 20 mM NaH_2_PO_4_/Na_2_HPO_4_ pH 7.4, 100 mM NaCl, 5 % glycerol and cOmplete™ EDTA-free protease inhibitor cocktail (Roche).

To determine the total membrane protein content, the membrane preparations were solubilized with 1% DDM and the total protein content was measured using Pierce™ 660 nm Protein Assay Reagent (Thermo Scientific) against the BSA standard curve (Thermo Scientific) in concentration range between 0.025 mg/mL and 2 mg/mL. The concentration of the sensor in the IMVs was determined from SDS-PAGE in-gel fluorescence with ImageQuant TL (Cytiva), using titrations of the purified sensors with known concentrations for the calibration.

### Characterization of the oligomeric state with SEC-MALS

The oligomeric state of the purified sensor constructs was analyzed by size exclusion chromatography coupled to multi-angle light scattering (SEC-MALS) using Superdex 200 Increase GL 10/300 column coupled to connected to miniDAWN TREOS II light scattering device and Optilab-TrEX Ri-detector (Wyatt Technology Corp.). The sensors were applied at 0.55 mg/mL concentrations in 10 mM NaH_2_PO_4_/Na_2_HPO_4_ pH 7.4, 50 mM NaCl and 0.05% DDM. Experiments with the soluble sensor construct lacking the transmembrane SecE domain were conducted at the same conditions in the buffer without DDM. The data analysis was performed with ASTRA 7.3.2 software (Wyatt Technology Corp.).

### Protein expression & characterization (crowding agents)

The protein crowding agents were expressed and purified as described elsewhere: mSA (Lim *et al*, 2011; Demonte *et al*, 2014), Strep^D4^ (Howarth *et al*, 2006), SecB (Fekkes *et al*, 1998), SecA^N^ and SecA^C^ (Kamel *et al*, 2022). As mSA was expressed as inclusion bodies and had to be refolded, its functionality was additionally analyzed by differential scanning fluorimetry (nanoDSF, Prometheus NT48). 1 µM mSA was optionally incubated with 10 µM biotin and the thermal denaturation of the protein was examined between 25 and 85°C (heating ramp 1°C/min) upon monitoring the intrinsic fluorescence at 330 and 350 nm, and the protein stabilization upon ligand binding was analyzed.

### Reconstitution of the crowding sensor into model membranes

Lipids were purchased in chloroform-solubilized form (Avanti Polar Lipids, Inc.) and were mixed together to obtain required lipid compositions. For PEG-based crowding experiments, liposomes composed of DOPC (63 mol %) and DOPG (27 mol %) were supplemented with 10 mol % of 1,2-dioleoyl-sn-glycero-3- phosphatidylethanolamine (DOPE) and 1,2-dioleoyl-sn-glycero-3-phosphoethanolamine-N-[methoxy(polyethylene glycol) -2000] (DOPE-PEG 2000) at various ratios. For protein-based crowding experiments, 20 mol % of anchor lipids, 1,2- dioleoyl-sn-glycero-3-[(N-(5-amino-1-carboxypentyl) iminodiacetic acid)succinyl] (18:1 DGS-NTA(Ni)) or 1,2-dioleoyl-sn-glycero-3-phosphoethanolamine-N-(cap biotinyl) (sodium salt) (18:1 Biotinyl Cap PE), were added to DOPC:DOPG mixture (53 mol% : 27 mol %) were used for titration experiments. Lipids were mixed in defined ratios, chloroform was removed via vacuum evaporation (rotary evaporator RV 8, IKA) while incubating the samples at 40 °C in a water bath. Formed lipid film was subsequently rehydrated and resuspended with 20 mM Tris-HCl pH 7.5 and 150 mM KCl to achieve final lipid concentration of 5 mM.

The liposome suspensions were extruded with the Mini-Extruder set (Avanti Polar Lipids, Inc.) via 0.2 µm polycarbonate membranes (Nuclepore, Whatman) and liposomes were swelled with 0.2 % DDM at 40°C for 15 min (Suppl. Figure 8). Unless other is indicated, the purified crowding sensors were added at the protein:lipid molar ratio of 1:3000 and incubated for 30 min on ice. Afterwards the samples were incubated with Bio-Beads SM-2 sorbent (Bio-Rad Laboratories) overnight on the rolling bank at 4°C to remove the detergent (Rigaud *et al*, 1997). Proteoliposomes with the reconstituted sensor were pelleted at 162000x*g* for 30 min (S120- AT3 rotor, Discovery M120 SE, Thermo Fisher/Sorvall) and then resuspended in 50 mM Tris-HCl pH 7.5 and 150 mM KCl to the final lipid concentration of 5 mM.

### Sensor reconstitution efficiency and topology analysis in liposomes

The reconstitution efficiency of the membrane-anchored crowding sensors was examined upon centrifugation in the sucrose density gradient. 50 µL of reconstituted proteoliposomes were mixed together with 60 % sucrose (w/v), 50 mM Tris-HCl pH 7.5 and 150 mM KCl to final sucrose concentration of 30 % in 200 µL, and loaded at the bottom of the centrifugation tube. 250 µL of 20 % sucrose solution and 50 µL of 5% sucrose solution were loaded on top, thus forming a step gradient of sucrose. The samples were centrifuged for 1 h at 29000x*g* (S120- AT3 rotor, Discovery M120 SE, Thermo Fisher/Sorvall) and then harvested from the bottom into 3 fractions (bottom” of 250 µL, “middle” 125 µL, and “top” of 125 µL). The presence of the sensor in each fraction was analyzed by SDS-PAGE: The intensity of fluorescent bands in SDS-PAGE was quantified (ImageQuant TL, Cytiva) and the relative amount of the reconstituted sensor was calculated by dividing band intensity of the individual fractions by the cumulative intensity of all fractions. Flotation experiments were carried independently at least two times for each sensor construct.

For studying the topology of the membrane-embedded sensors, DOPC:DOPG liposomes were incubated with 0.2 % DDM or 0.5 % Triton X-100, and *α*H-SecE or (GSG)_6_-SecE sensors were reconstituted as described above. Formed proteoliposomes were mixed with either 42 µM trypsin (from porcine pancreas, Sigma-Aldrich) or 17 µM proteinase K (Thermo Fisher Scientific). Detergent-solubilized sensors were equally incubated with proteases and served as controls in this experiment. The proteolysis reaction proceeded for 2 h at 22 °C, then the samples were incubated for 5 min at 90 °C to inactivate the proteases and were analyzed by SDS-PAGE.

### Fluorescence spectroscopy

Purified and optionally reconstituted sensors were diluted in 20 mM Tris-HCl pH 7.4 and 150 KCl, and the emission spectrums of the probes were recorded on either Fluorolog-3 or FluoroMax-Plus (Horiba™ Scientific). The excitation wavelength was set to 420 nm, slit width 5 nm, so only the donor fluorophore mCerulean was excited, and the fluorescence emission spectra were recorded in the range of 435 to 620 nm, where the emission of mCerulean (donor) was measured at 475 nm, and mCitrine (acceptor) at 525 nm. Dilution series of PEG 6000 as a soluble crowder were prepared in 20 mM Tris-HCl pH 7.4 and 150 mM KCl based on 50 % stock solution (w/v). For measurements that included the detergent-solubilized sensors, 0.05 % DDM was additionally supplemented. To induce protein crowding at the liposome surface, crowders were titrated stepwise to the liposomes with reconstituted sensors until the crowder/ligand-lipid ratio of 1.1 was reached. To probe crowding in IMVs, Strep^D4^ and SecA were added to vesicle suspension in concentrations of 13 µM for αH-SecE and 8 µM for (GSG)_6_-SecE samples. For all the samples the background spectrum of the corresponding buffer or crowder solution was subtracted.

### Super-resolution structured illumination microscopy

Cells transformed with pBAD-based plasmids containing genes for either (GSG)_6_-SecE or *α*H- SecE sensors were grown as described earlier. Additional cell culture with *α*H-SecE sensor was prepared as a control and was not induced with arabinose. The harvested cells were resuspended in PBS and the OD_600_ was adjusted to 1.2. Cover glasses for the microscopy were cleaned with 70% ethanol and coated by 0.1% (w/v) poly-L-lysine solution. Next, the cover glasses were placed into 12-well plates with 1 mL PBS and 5 µL of bacterial cell suspension and centrifuged at 1500 rpm (ROTOR) for 15 min at 4°C. The supernatant was removed and the attached cells were washed with 1 mL of fresh PBS. Structured illumination microscopy was performed using the Zeiss ELYRA PS.1 microscope system (Zeiss Microscopy GmbH, Oberkochen, Germany) equipped with a Plan-Apochromat 63x/1.4 oil immersion objective lens. For excitation of the sensors a 488 nm diode laser was used at 1,5- 2,5% emission intensity. Signals were detected by a front illuminated Andor iXon3 DU-885K camera, a BP 495-575 + LP 750 emission filter, exposure time of 100 ms and an EMCCD gain of 100-200. Individual stacks of 256x256 px (pixel) and a Z-axis interval of 110nm were acquired at 5 42µm SIM-grid rotations and with no averaging. Each acquired z-stack was processed internally with the ZEN black SIM feature with the same 3D signal-to-noise filter of -3,3 for all data.

## Acknowledgements

The work was supported by the German Research Foundation (Deutsche Forschungsgemeinschaft, DFG) via the Research Grant Ke1879/3 and the Collaborative Research Center 1208 “Identity and Dynamics of Membrane Systems”. We thank Prof. Arnold J. Boersma (University of Utrecht), Dr. Jens Reiners and Dr. Jakub Kubiak for technical advice and discussions along the project.

## References

1. Armstrong JK, Wenby RB, Meiselman HJ & Fisher TC (2004) The hydrodynamic radii of macromolecules and their effect on red blood cell aggregation. Biophys J 87: 4259– 4270

2. Aumiller WM, Davis BW, Hatzakis E & Keating CD (2014) Interactions of macromolecular crowding agents and cosolutes with small-molecule substrates: Effect on horseradish peroxidase activity with two different substrates. J Phys Chem B 118: 10624–10632

3. Bai J, Liu M, Pielak GJ & Li C (2017) Macromolecular and Small Molecular Crowding Have Similar Effects on alpha-Synuclein Structure. Chemphyschem 18: 55–58

4. Berg B van den, Ellis RJ & M.Dobson C (1999) Effects of macromolecular crowding on protein folding and aggregation. EMBO J 18: 6927–6933

5. Boersma AJ, Zuhorn IS & Poolman B (2015) A sensor for quantification of macromolecular crowding in living cells. Nat Methods 12: 227–229

6. Bohrmann B, Haider M & Kellenberger E (1993) Concentration evaluation of chromatin in unstained resin-embedded sections by means of low-dose ratio-contrast imaging in STEM. Ultramicroscopy 49: 235–251

7. Bokvist M & Grobner G (2007) Misfolding of Amyloidogenic Protein at Membrane Surfaces: Impact of Macromolecular Crowding. J Am Chem Soc 129: 14848–14849

8. Breyton C, Haase W, Rapoport TA, Kühlbrandt W & Collinson I (2002) Three-dimensional structure of the bacterial protein-translocation complex SecYEG. Nature 418: 662–665

9. Chen L, Novicky L, Merzlyakov M, Hristov T & Hristova K (2010) Measuring the energetics of membrane protein dimerization in mammalian membranes. J Am Chem Soc 132: 3628– 3635

10. Collot M, Pfister S & Klymchenko AS (2022) Advanced functional fluorescent probes for cell plasma membranes. Curr Opin Chem Biol 69: 102161

11. Demonte D, Dundas CM & Park S (2014) Expression and purification of soluble monomeric streptavidin in Escherichia coli. Appl Microbiol Biotechnol 98: 6285–6295

12. Dupuy AD & Engelman DM (2008) Protein area occupancy at the center of the red blood cell membrane. Proc Natl Acad Sci U S A 105: 2848–2852

13. Elowitz MB, Surette MG, Wolf PE, Stock JB & Leibler S (1999) Protein mobility in the cytoplasm of Escherichia coli. J Bacteriol 181: 197–203

14. Engelman DM & Steitz TA (1981) The spontaneous insertion of proteins into and across membranes: The helical hairpin hypothesis. Cell 23: 411–422

15. Fekkes P, De Wit JG, Van Der Wolk JPW, Kimsey HH, Kumamoto CA & Driessen AJM (1998) Preprotein transfer to the Escherichia coli translocase requires the co-operative binding of SecB and the signal sequence to SecA. Mol Microbiol 29: 1179–1190

16. Fotiadis D, Liang Y, Filipek S, Saperstein DA, Engel A & Palczewski K (2003) Rhodopsin dimers in native disc membranes. Nature 421: 127–128

17. Geertsma ER, Nik Mahmood NAB, Schuurman-Wolters GK & Poolman B (2008) Membrane reconstitution of ABC transporters and assays of translocator function. Nat Protoc 3: 256–266

18. Guigas G & Weiss M (2016) Effects of protein crowding on membrane systems. Biochim Biophys Acta - Biomembr 1858: 2441–2450

19. Guseman AJ, Perez Goncalves GM, Speer SL, Young GB & Pielak GJ (2018) Protein shape modulates crowding effects. Proc Natl Acad Sci U S A 115: 10965–10970

20. Hills RD & McGlinchey N (2016) Model parameters for simulation of physiological lipids. J Comput Chem 37: 1112–1118

21. Houser JR, Hayden CC, Thirumalai D & Stachowiak JC (2020) A Förster Resonance Energy Transfer-Based Sensor of Steric Pressure on Membrane Surfaces. J Am Chem Soc 142: 20796–20805

22. Howarth M, Chinnapen DJF, Gerrow K, Dorrestein PC, Grandy MR, Kelleher NL, El-Husseini A & Ting AY (2006) A monovalent streptavidin with a single femtomolar biotin binding site. Nat Methods 3: 267–273

23. Janovjak H, Struckmeier J, Hubain M, Kedrov A, Kessler M & Müller DJ (2004) Probing the energy landscape of the membrane protein bacteriorhodopsin. Structure 12

24. Kamel M, Löwe M, Schott-Verdugo S, Gohlke H & Kedrov A (2022) Unsaturated fatty acids augment protein transport via the SecA:SecYEG translocon. FEBS J 289: 140–162

25. Karagöz GE, Acosta-Alvear D & Walter P (2019) The Unfolded Protein Response: Detecting and Responding to Fluctuations in the Protein-Folding Capacity of the Endoplasmic Reticulum. Cold Spring Harb Perspect Biol: a033886

26. Kater L, Frieg B, Berninghausen O, Gohlke H, Beckmann R & Kedrov A (2019) Partially inserted nascent chain unzips the lateral gate of the Sec translocon. EMBO Rep 20: e48191

27. Kedrov A, Ziegler C, Janovjak H, Kühlbrandt W & Müller DJ (2004) Controlled unfolding and refolding of a single sodium-proton antiporter using atomic force microscopy. J Mol Biol 340

28. Kirchhoff H (2008) Molecular crowding and order in photosynthetic membranes. Trends Plant Sci 13: 201–207

29. Kuznetsova IM, Turoverov KK & Uversky VN (2014) What Macromolecular Crowding Can Do to a Protein. Int J Mol Sci 15: 23090–23140

30. Kuznetsova IM, Zaslavsky BY, Breydo L, Turoverov KK & Uversky VN (2015) Beyond the excluded volume effects: Mechanistic complexity of the crowded milieu. Molecules 20: 1377–1409

31. Lambert TJ (2019) FPbase: a community-editable fluorescent protein database. Nat Methods 16: 277–278

32. Lim KH, Huang H, Pralle A & Park S (2011) Engineered streptavidin monomer and dimer with improved stability and function. Biochemistry 50: 8682–8691

33. Liu B, Åberg C, van Eerden FJ, Marrink SJ, Poolman B & Boersma AJ (2017) Design and Properties of Genetically Encoded Probes for Sensing Macromolecular Crowding. Biophys J 112: 1929–1939

34. Liu LN & Scheuring S (2013) Investigation of photosynthetic membrane structure using atomic force microscopy. Trends Plant Sci 18: 277–286

35. Löwe M, Kalacheva M, Boersma AJ & Kedrov A (2020) The more the merrier: effects of macromolecular crowding on the structure and dynamics of biological membranes. FEBS J 287: 5039–5067

36. Marsh D, Bartucci R & Sportelli L (2003) Lipid membranes with grafted polymers: Physicochemical aspects. Biochim Biophys Acta - Biomembr 1615: 33–59

37. Minton AP & Wilf J (1981) Effect of Macromolecular Crowding upon the Structure and Function of an Enzyme: Glyceraldehyde-3-phosphate Dehydrogenase. Biochemistry 20: 4821–4826

38. Mouton SN, Thaller DJ, Crane MM, Rempel IL, Terpstra O, Steen A, Kaeberlein M, Lusk CP, Boersma AJ & Veenhoff LM (2020) A physicochemical perspective of aging from single- cell analysis of ph, macromolecular and organellar crowding in yeast. Elife 9: 1–42

39. Nawrocki G, Wang PH, Yu I, Sugita Y & Feig M (2017) Slow-Down in Diffusion in Crowded Protein Solutions Correlates with Transient Cluster Formation. J Phys Chem B 121: 11072–11084

40. Nemzer LR, Flanders BN, Schmit JD, Chakrabarti A & Sorensen CM (2013) Ethanol shock and lysozyme aggregation. Soft Matter 9: 2187–2196

41. Niroomand H, Venkatesan GA, Sarles SA, Mukherjee D & Khomami B (2016) Lipid- Detergent Phase Transitions During Detergent-Mediated Liposome Solubilization. J Membr Biol 249: 523–538

42. Prévost C, Sharp ME, Kory N, Lin Q, Voth GA, Farese RVJ & Walther TC (2018) Mechanism and Determinants of Amphipathic Helix-Containing Protein Targeting to Lipid Droplets. Dev Cell 44: 73–86

43. Raghunath G & Dyer RB (2019) Kinetics of Histidine-Tagged Protein Association to Nickel- Decorated Liposome Surfaces. Langmuir 35: 12550–12561

44. Rigaud JL, Mosser G, Lacapere JJ, Olofsson A, Levy D & Ranck JL (1997) Bio-Beads: An efficient strategy for two-dimensional crystallization of membrane proteins. J Struct Biol 118: 226–235

45. Rivas G & Minton AP (2018) Toward an understanding of biochemical equilibria within living cells. Biophys Rev 10: 241–253

46. Rohwer JM, Postma PW, Kholodenko BN & Westerhoff H V. (1998) Implications of macromolecular crowding for signal transduction and metabolite channeling. Proc Natl Acad Sci U S A 95: 10547–10552

47. Roussel G & White SH (2020) The SecA ATPase motor protein binds to Escherichia coli liposomes only as monomers. Biochim Biophys Acta - Biomembr 1862: 183358

48. Sarkar M, Lu J & Pielak GJ (2014) Protein-Crowder Charge and Protein Stability. Biochemistry 53: 1601–1606

49. Sezgin E (2017) Super-resolution optical microscopy for studying membrane structure and dynamics. J Phys Condens Matter 29

50. Sezgin E, Levental I, Mayor S & Eggeling C (2017) The mystery of membrane organization: Composition, regulation and roles of lipid rafts. Nat Rev Mol Cell Biol 18: 361–374

51. Sharpe HJ, Stevens TJ & Munro S (2010) A Comprehensive Comparison of Transmembrane Domains Reveals Organelle-Specific Properties. Cell 142: 158–169

52. Smith VF, Schwartz BL, Randall LL & Smith RD (1996) Electrospray mass spectrometric investigation of the chaperone SecB. Protein Sci 5: 488–494

53. Srere PA (1980) The infrastructure of the mitochondrial matrix. Trends Biochem Sci 5: 120– 121

54. Strop P & Brunger AT (2005) Refractive index-based determination of detergent concentration and its application to the study of membrane proteins. Protein Sci 14: 2207–2211

55. Wang H, Chan SH, Dey S, Castello-serrano I, Rosen MK, Ditlev JA, Levental KR & Levental I (2023) Coupling of protein condensates to ordered lipid domains determines functional membrane organization.

56. Zakeri B, Fierer JO, Celik E, Chittock EC, Schwarz-Linek U, Moy VT & Howarth M (2012) Peptide tag forming a rapid covalent bond to a protein, through engineering a bacterial adhesin. Proc Natl Acad Sci 109: E690–E697

57. Zimmerman SB & Pheiffer BH (1983) Macromolecular crowding allows blunt-end ligation by DNA ligases from rat liver or Escherichia coli. Proc Natl Acad Sci U S A 80: 5852–5856

58. Zimmerman SB & Trach SO (1991) Estimation of macromolecule concentrations and excluded volume effects for the cytoplasm of Escherichia coli. J Mol Biol 222: 599–620

