## Supplemental data for "Probing macromolecular crowding at the lipid membrane interface with genetically-encoded sensors"

**Supplemental figures**

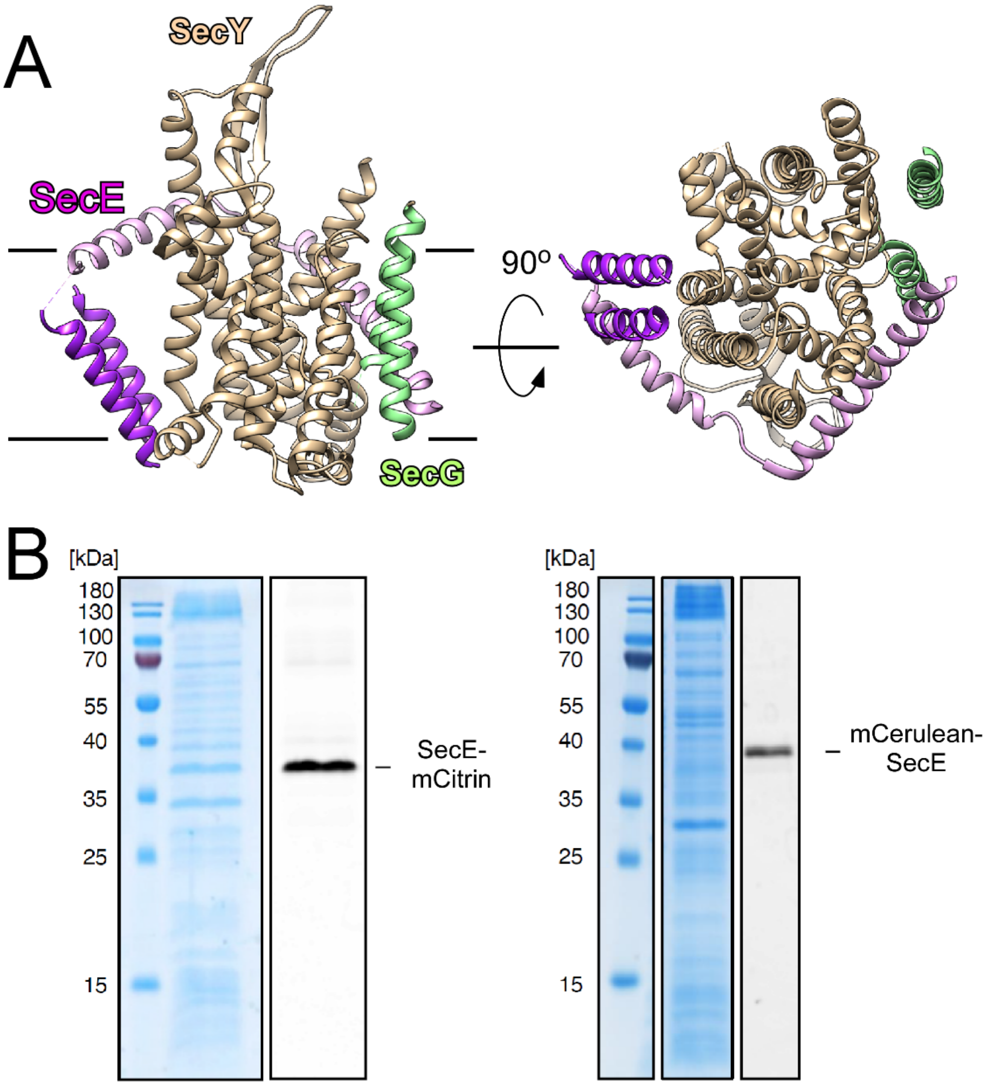


**Supplemental Figure 1. Transmembrane anchor for the crowding sensors.** (A) SecYEG translocon in the lipid membrane (PDB: 6R7L), view in the membrane plane (left) and from the periplasm (right). The membrane plane is shown by red lines, as resolved by cryo-EM . Individual subunits are indicated and color-coded. SecE TMHs 1-2 used for anchoring the crowding sensors are shown in dark purple. The connecting periplasmic loop of four amino acids is not resolved in the cryo-EM structure. (B) SDS-PAGE of crude membrane extracts containing SecE-mCitrine and mCerulean-SecE fusion proteins. Left: Coomassie stained gels; right: In-gel fluorescence.

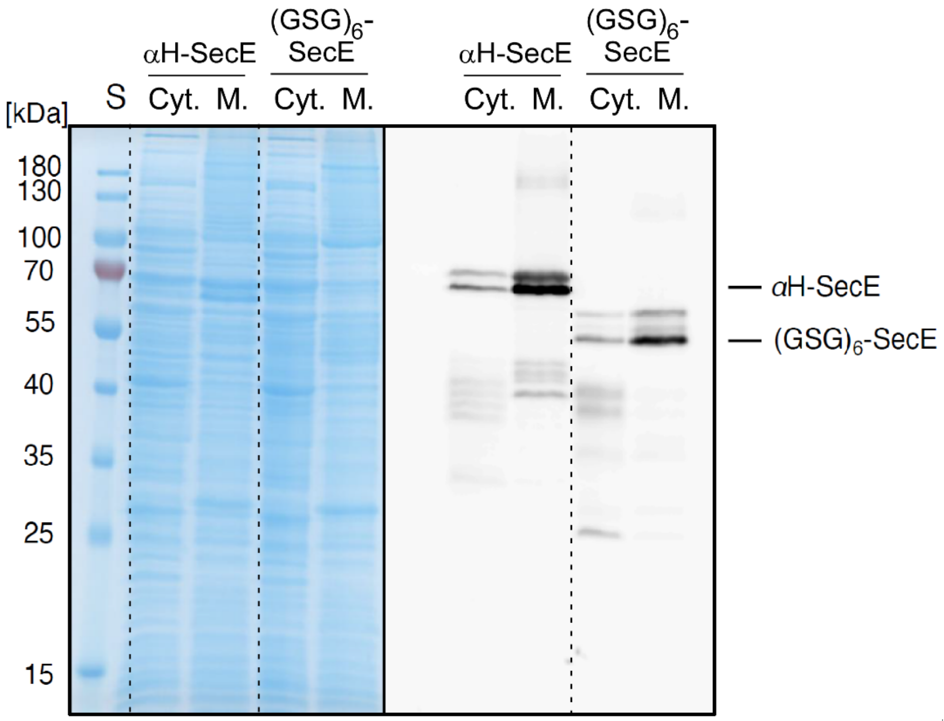


**Supplemental Figure 2. Expression of the crowding sensors.** SDS-PAGE (left) and in-gel fluorescence (right) of the cytoplasmic (“Cyt.”) and crude membrane (“M.”) fractions of the cells expressing the indicated crowding sensor.

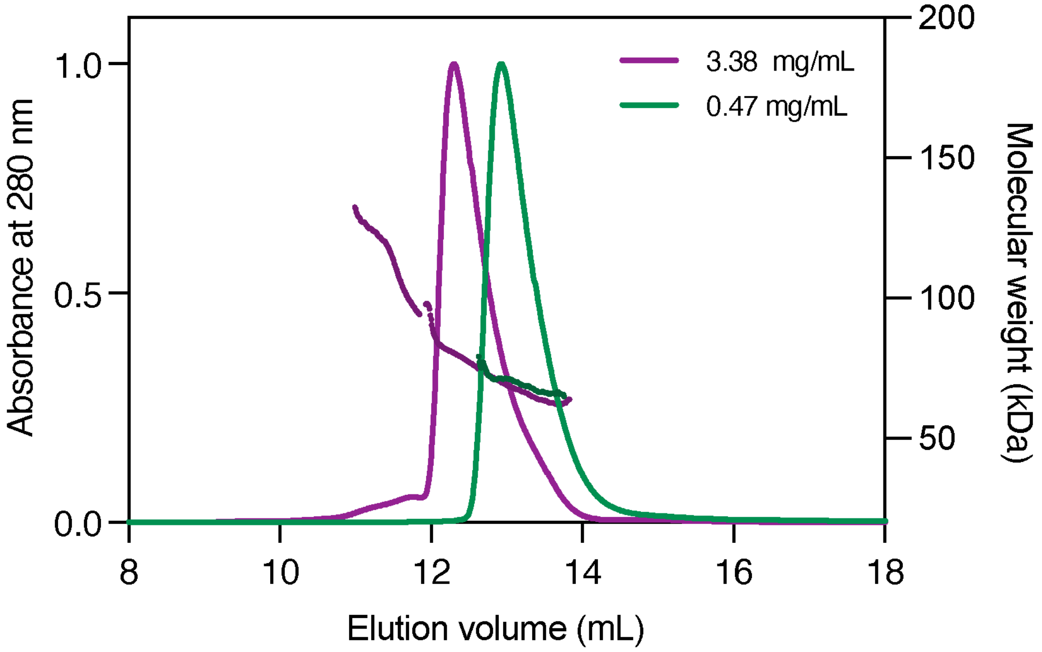


**Supplemental Figure 3. Soluble sensor undergoes concentration-dependent oligomerization.** SEC-MALS profiles of the membrane anchor-free sensor at 0.47 and 3.38 mg/mL concentrations revels a shift towards higher molecular mass at the elevated concentration. For the sample injected with 0.47 mg/mL (green), only one peak in MALS with 70.4 ± 1.5 kDa was measured. For the elevated concentration (purple), several peaks were detected at 269 ± 16 kDa, 110 ± 1 kDa and 75 ± 1 kDa, thus showing a concentration-dependent oligomerization of the sensor in absence of the hydrophobic anchor.

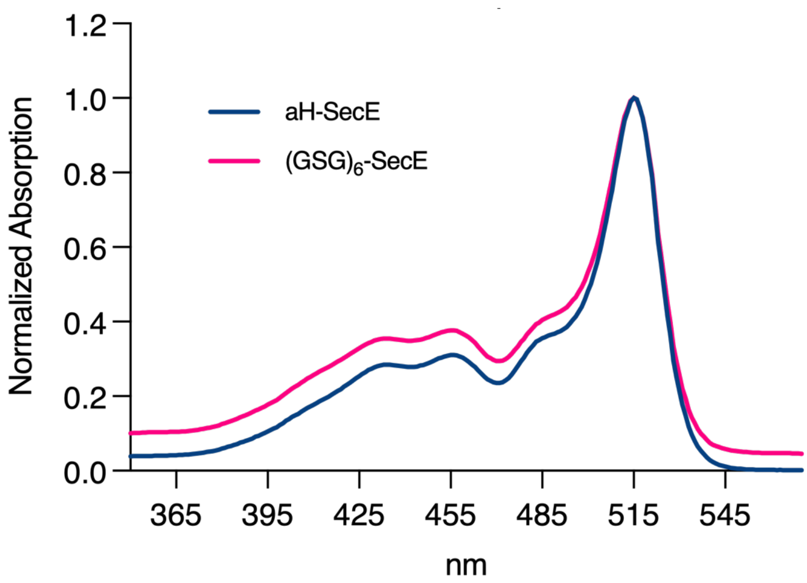


**Supplemental Figure 4. Absorption spectra of the purified crowding sensors.** The characteristic absorption spectra confirm presence of two fluorescent proteins with specific peaks for mCerulean (~430 nm) and mCitrine (515 nm).

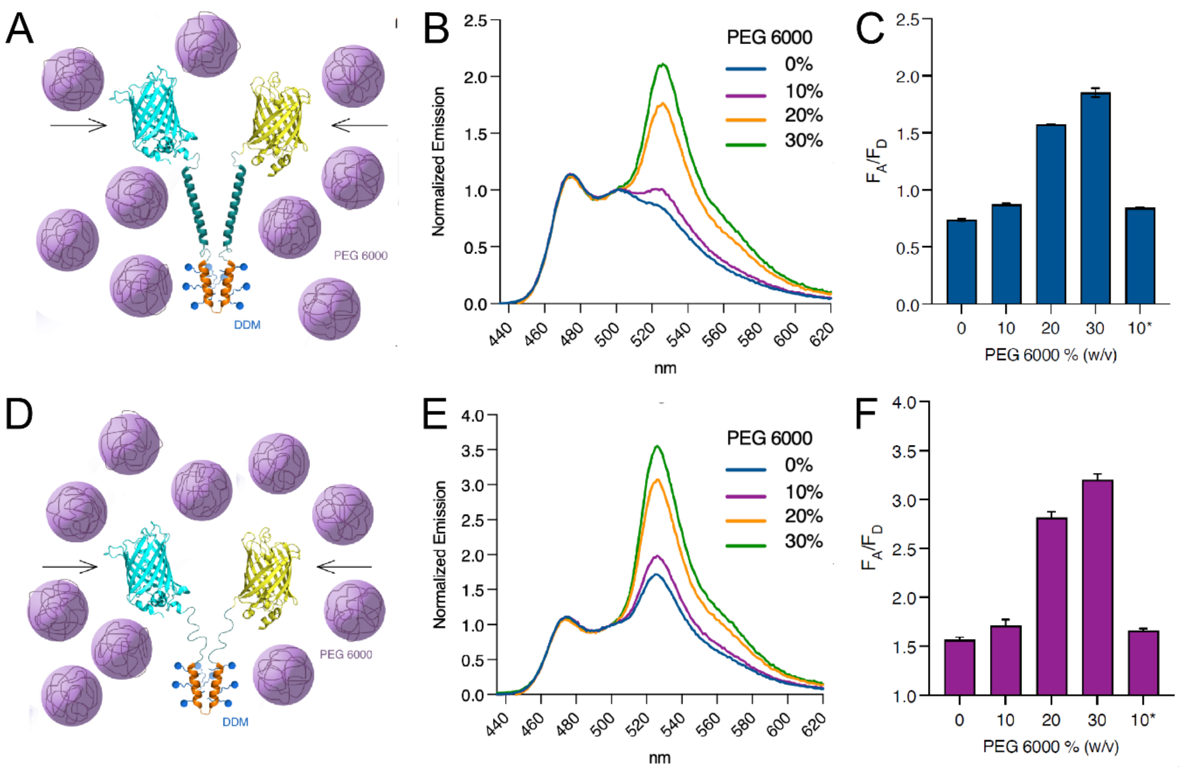


**Supplemental Figure 5. Sensitivity of the detergent-solubilized sensors to the polymer-induced crowding.** (A) Scheme of the $\alpha$H-SecE sensor in detergent micelle upon compaction induced by the polymer in solution. (B) Fluorescence emission spectra of $\alpha$H-SecE sensor in presence of PEG 6000 at indicated concentrations (w/v). (C) Corresponding FRET efficiencies of $\alpha$H-SecE sensor (mean ± SD, n = 2). Sample "10%*" correspond to two-fold dilution of 20% PEG 6000 for testing the reversibility of the sensor compaction. (D-F) Same as (A-C), for the detergent-solubilized (GSG)$_6$-SecE sensor.

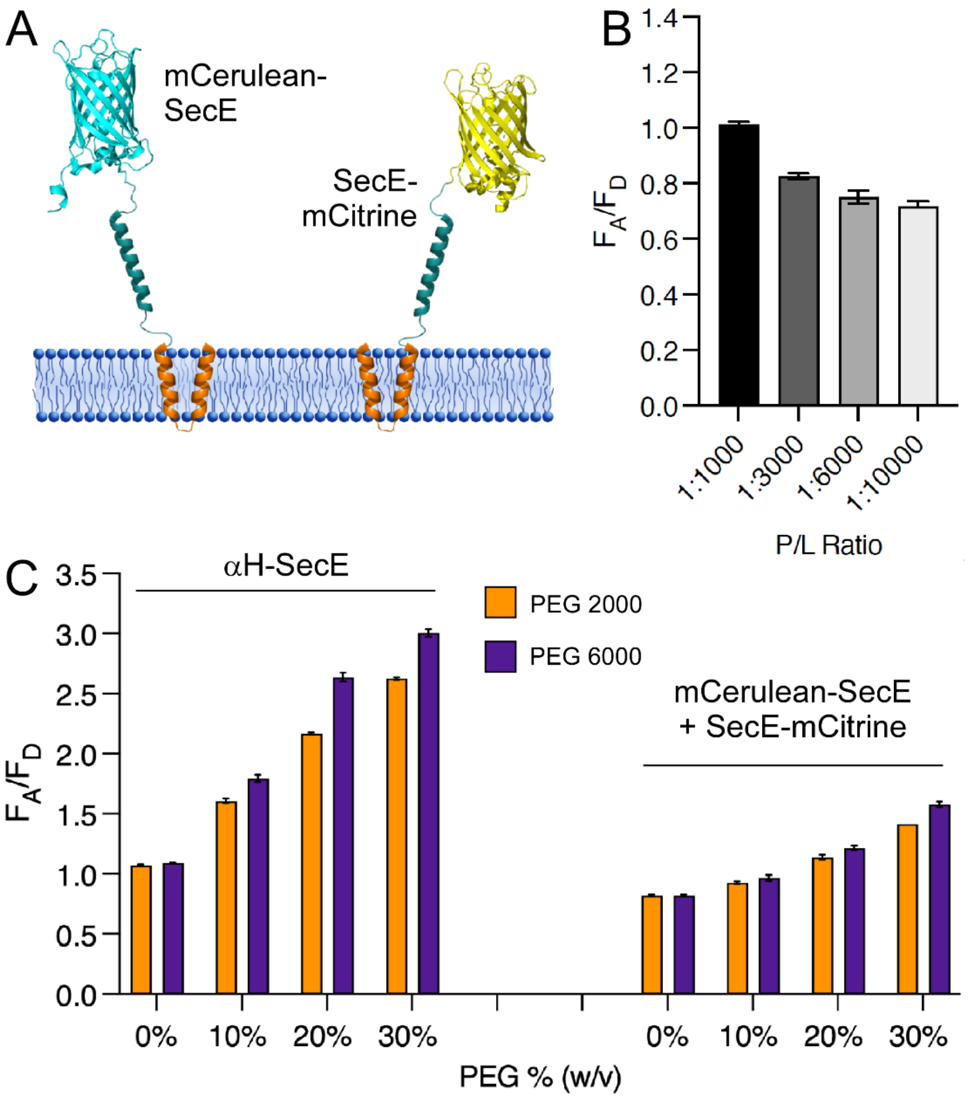


**Supplemental Figure 6. Individual membrane-anchored fluorophores manifest weak dependence of FRET on macromolecular crowding**. (A) Scheme of co-reconstituted fluorophores fused with individual membrane anchors. (B) FRET efficiency is dependent on the protein-to-lipid ratio used for reconstitution. Elevated protein density facilitates intermolecular FRET. (C) FRET efficiency of the crowding sensor αH-SecE vs. individual co-reconstituted fluorophores at P/L ratio of 1:3,000 in response to soluble crowders PEG 2000 and PEG 6000.

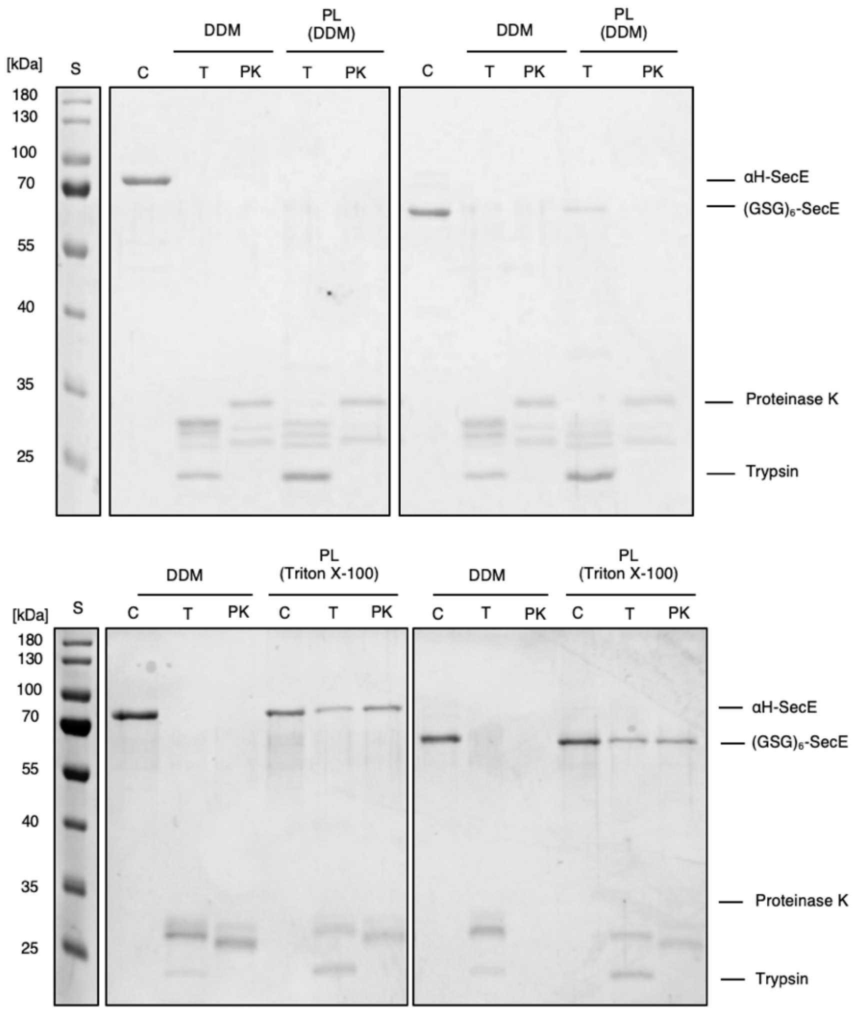


**Supplemental Figure 7. Topology determination of the liposome-reconstituted sensor using limited proteolysis.** Topology determination of the liposome-reconstituted sensors via limited proteolysis by trypsin, “T”, or proteinase K, “PK”. “DDM”, detergent-solubilized sensors, “PL (DDM)”, sensors in proteoliposomes reconstituted using DDM. (B) Same as (A), but using Triton X-100 for the reconstitution.

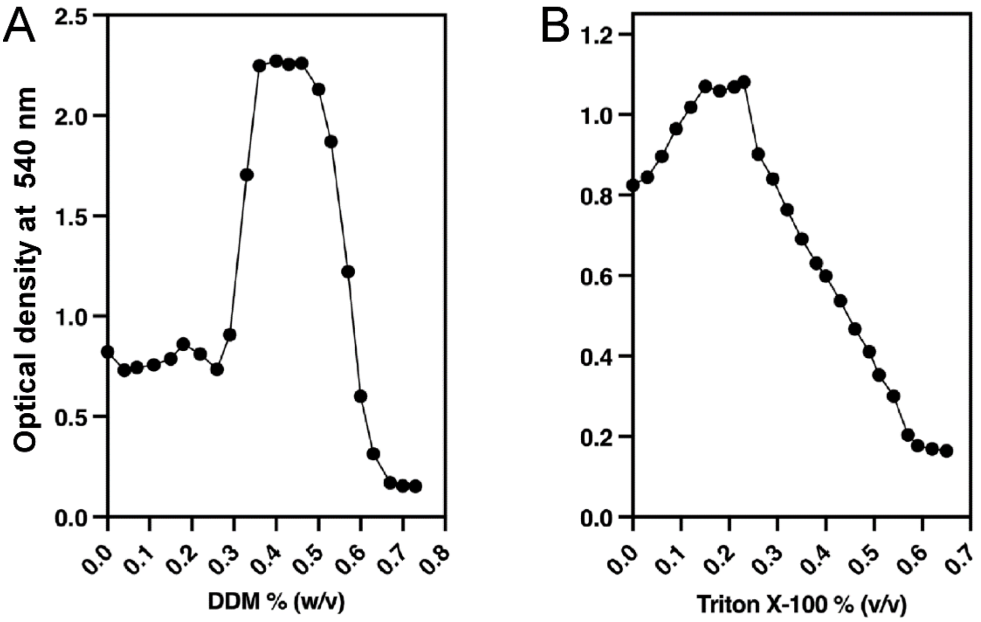


**Supplemental Figure 8**. **Destabilization of liposomes prior reconstitution of the crowding sensors.** Optical density for DOPC:DOPG (70:30 mol%) liposomes in presence of DDM (A) and Triton X-100 (B). The liposomes were titrated with detergents and the optical density at 540 nm was recorded after each step until the suspension was completely solubilized. At the beginning of the titrations the liposome swelling in observed for both detergents resulting in increase of the optical density, followed by disintegration/solubilization of liposomes and decrease in the optical density.

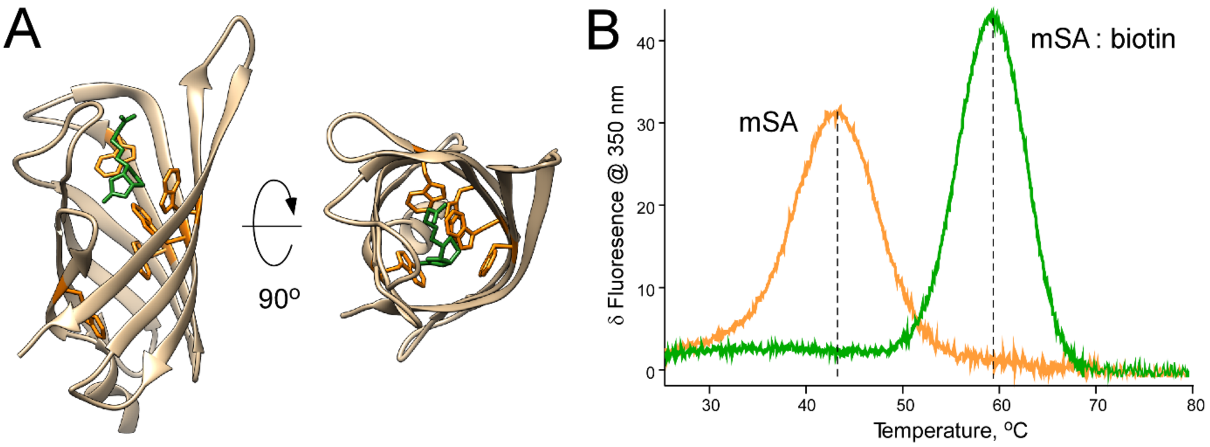


**Supplemental Figure 9. Functional test of the refolded monomeric streptavidin (mSA).** (A) Structure of mSA with bound biotin molecule (PDB ID: 4JNJ). Tryptophan residues are shown in orange, biotin in green. (B) Differential scanning fluorometry of mSA and mSA:biotin complex. Biotin binding induces prominent thermodynamic stabilization of mSA, as the thermal denaturation point shifts from 43 °C to 59 °C. Complete shift to the higher denaturation temperature indicates that all mSA molecules were competent for the ligand binding.

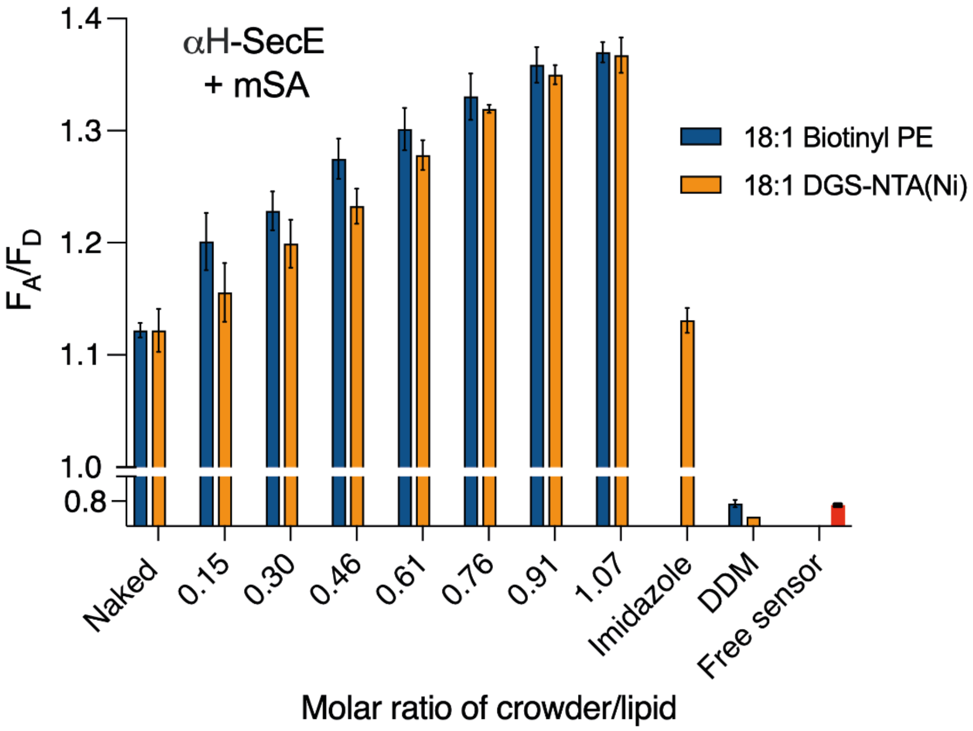


**Supplemental Figure 10. mSA-induced crowding at the membrane interface is detected by the sensor.** Two approaches for binding mSA to the membrane surface, either via biotinyl cap PE lipids or DGS-NTA lipids resulted in nearly identical response of the reconstituted sensor αH-SecE.

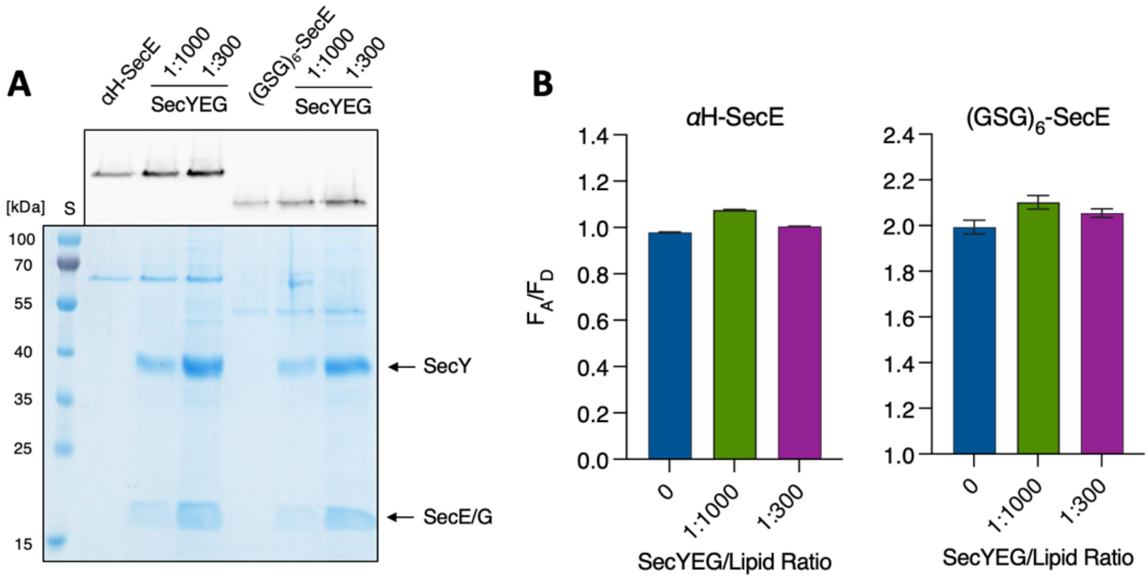


**Supplemental Figure 11. Interfacial sensors do not respond to the crowding within the membrane.** (A) Protein content of the proteoliposomes with either sensors alone or also SecYEG translocon added at the indicated protein-to-lipid molar ratios. (B) FRET efficiencies of both sensors in each type of proteoliposomes.

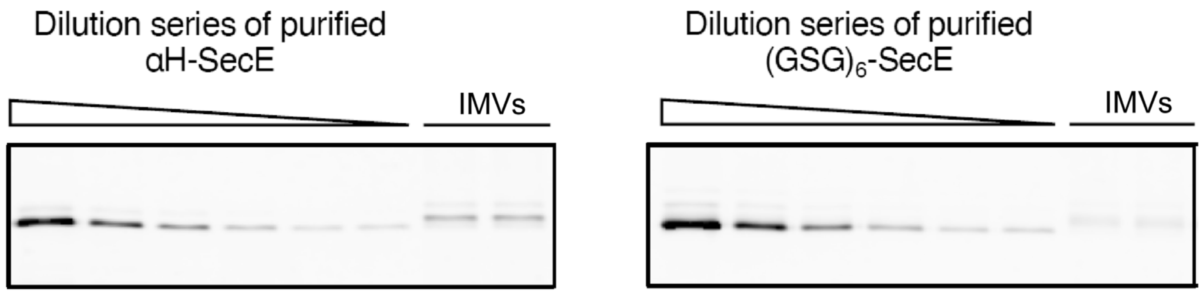


**Supplemental Figure 12. Determination of the sensor abundance in the bacterial membrane.** In-gel fluorescence images of SDS-PAGE show IMVs containing the sensors loaded next to the serial dilutions of the purified sensors of known concentrations (range 166 μg/mL to 5 μg/mL).

**$\alpha$H-SecE sensor amino acid sequence:**

MHHHHHHLEVLFQGPGVSKGEELFTGVVPILVELDGDVNGHKFSVSGEGEGDATYGKLTLKFICTTGKLPVPWPTLVTTLSWGVQCFARYPDHMKQHDFFKSAMPEGYVQERTIFFKDDGNYKTRAEVKFEGDTLVNRIELKGIDFKEDGNILGHKLEYNAIHGNVYITADKQKNGIKANFGLNCNIEDGSVQLADHYQQNTPIGDGPVLLPDNHYLSTQSKLSKDPNEKRDHMVLLEFVTAAGITLGMDELYKGSGGSGGSGGSGGSGGSGAEAAAKEAAAKEAAAKEAAAKEAAAKEAAAKAGSGGSGGSGGSGANTEAQGSGRGLEAMKWVVVVALLLVAIVGNYLYRDIMLPLRALAVVILIAAAGGVALLTTKGKGSGGSGGSGGSGAEAAAKEAAAKEAAAKEAAAKEAAAKEAAAKAGSGGSGGSGGSGGSGGSMVSKGEELFTGVVPILVELDGDVNGHKFSVSGEGEGDATYGKLTLKFICTTGKLPVPWPTLVTTFGYGLMCFARYPDHMKQHDFFKSAMPEGYVQERTIFFKDDGNYKTRAEVKFEGDTLVNRIELKGIDFKEDGNILGHKLEYNYNSHNVYIMADKQKNGIKVNFKIRHNIEDGSVQLADHYQQNTPIGDGPVLLPDNHYLSYQSALSKDPNEKRDHMVLLEFVTAAGITLGMDELYK

**(GSG)$_6$-SecE sensor amino acid sequence:**

MHHHHHHLEVLFQGPGVSKGEELFTGVVPILVELDGDVNGHKFSVSGEGEGDATYGKLTLKFICTTGKLPVPWPTLVTTLSWGVQCFARYPDHMKQHDFFKSAMPEGYVQERTIFFKDDGNYKTRAEVKFEGDTLVNRIELKGIDFKEDGNILGHKLEYNAIHGNVYITADKQKNGIKANFGLNCNIEDGSVQLADHYQQNTPIGDGPVLLPDNHYLSTQSKLSKDPNEKRDHMVLLEFVTAAGITLGMDELYKGSGGSGGSGGSGGSGGSGSANTEAQGSGRGLEAMKWVVVVALLLVAIVGNYLYRDIMLPLRALAVVILIAAAGGVALLTTKGKGSGGSGGSGGSGGSGGSMVSKGEELFTGVVPILVELDGDVNGHKFSVSGEGEGDATYGKLTLKFICTTGKLPVPWPTLVTTFGYGLMCFARYPDHMKQHDFFKSAMPEGYVQERTIFFKDDGNYKTRAEVKFEGDTLVNRIELKGIDFKEDGNILGHKLEYNYNSHNVYIMADKQKNGIKVNFKIRHNIEDGSVQLADHYQQNTPIGDGPVLLPDNHYLSYQSALSKDPNEKRDHMVLLEFVTAAGITLGMDELYK

mCerulean is highlighted in blue, mCitrine in orange, SecE in red, $\alpha$-helices of the linker region in green

1
